# SIRT7 regulates NUCKS1 chromatin binding to elicit metabolic and inflammatory gene expression in senescence and liver aging

**DOI:** 10.1101/2024.02.05.578810

**Authors:** Khoa Tran, Michael Gilbert, Berta N. Vazquez, Alessandro Ianni, Benjamin A. Garcia, Alejandro Vaquero, Shelley Berger

## Abstract

Sirtuins, a class of highly conserved histone/protein deacetylases, are heavily implicated in senescence and aging. The regulation of sirtuin proteins is tightly controlled both transcriptionally and translationally and via localization within the cell. While Sirtiun proteins are implicated with aging, how their levels are regulated during aging across cell types and eliciting tissue specific age-related cellular changes is unclear. Here, we demonstrate that SIRT7 is targeted for degradation during senescence and liver aging. To uncover the significance of SIRT7 loss, we performed proteomics analysis and identified a new SIRT7 interactor, the HMG box protein NUCKS1. We found that the NUCKS1 transcription factor is recruited onto chromatin during senescence and this is mediated by SIRT7 loss. Further, depletion of NUCKS1 delayed senescence upon DNA damage leading to reduction of inflammatory gene expression. Examination of NUCKS1 transcriptional regulation during senescence revealed gene targets of transcription factors NFKB1, RELA, and CEBPβ. Consistently, in both Sirt7 KO mouse liver and in naturally aged livers, Nucks1 was recruited to chromatin. Further, Nucks1 was bound at promoters and enhancers of age-related genes, including transcription factor Rela, and, moreover, these bound sites had increased accessibility during aging. Overall, our results uncover NUCKS1 as a novel interactor of SIRT7, and show that loss of SIRT7 during senescence and liver aging promotes NUCKS1 chromatin binding to regulate metabolic and inflammatory genes.

## INTRODUCTION

Aging is a highly complex cellular process characterized by damage drivers (e.g. genomic instability, epigenetic alteration), altered responses to damage (e.g. mitochondrial dysfunction, loss of nutrient-sensing), and downstream phenotypic consequences (e.g. cellular senescence, immune system decline)^1^. Aging is a critical risk factor for numerous diseases including cancer, cardiovascular disease, and neurological disorders^2,3^. While aging plays an essential role in these disorders, the underlying biology of aging and its contribution to these age-related diseases remain highly studied with key mechanisms still to be revealed.

Sirtuins are a family of NAD+-dependent deacetylases and ADP-ribosyltransferases that are heavily implicated in aging and senescence^4,5^, and individual sirtuins regulate lifespan through distinct mechanisms^5^. Sirtuins are distinctly localized in different compartments within the cell, for example, SIRT1, SIRT6, and SIRT7 localize within the nucleus^6^. Sirtuins are protein deacetylases and one focus of nuclear sirtuins is deacetylation of histones in chromatin. Previously, our lab demonstrated that *S. cerevisiae* SIR2 (homologous to human SIRT1-7) is proteolyzed during aging, resulting in accumulation of H4K16ac^7^. Further, our lab showed that mammalian SIRT1 is degraded via nuclear-initiated autophagy during cellular senescence and during aging of specific tissues^8^. Whether other sirtuin family members are post-transcriptionally regulated during senescence and aging, and the functional consequences during tissue aging of sirtuin post-transcriptional regulation remains unexplored.

SIRT7 is one of the least characterized Sirtuins, however, one important aspect is its role in senescence and aging^9,10^. Deletion of SIRT7 in mice leads to progeria-like phenotypes with altered liver function^11,12^. Further, depletion of SIRT7 in human fibroblasts leads to senescence due to increased genomic instability at ribosomal DNA defective maintenance of rDNA heterochromatin^13^. The depletion of SIRT7 in human mesenchymal stem cells accelerates senescence and leads to deregulation of heterochromatin and upregulation of LINE-1 elements to initiate the innate immune response^14,15^. While SIRT7 resgulates histone acetylation, SIRT7 also deacetylates non-histone protein targets, including p53^16^, DDB1^17^, CDK9^18^, PCAF^19^, CRY1^20^, NPM1^21^, HMGB1^22^, and has mono-ADP ribosylation activity including self-ribosylation^23^. SIRT7-dependent deacetylation of non-histone targets is critical for mediating the stress response; SIRT7 regulates p53 activity by directly deacetylating the protein itself^16^ or regulators of p53^19,21,24^. Notably, while non-histone proteins of SIRT7 are targeted in cellular senescence, the functional impacts during tissue aging of altered acetylation is unclear.

Here, we report that SIRT7 protein is targeted for degradation during cellular senescence and is reduced in aged mouse liver, with SIRT7 knockout in young liver leading to aging-like gene expression. We uncover interaction of SIRT7 with the HMG box containing transcription factor NUCKS1—which has a documented role in cell cycle^25^, inflammatory signaling^26^ and DNA damage^27,28^, all processes implicated in senescence—and show that SIRT7 loss during senescence or in liver promotes NUCKS1 recruitment onto chromatin and thereby alters relevant gene expression. Overall, our findings in senescence and aging reveal an unanticipated mechanism of SIRT7 through regulation of the NUCKS1 transcription factor.

## RESULTS

### SIRT7 is targeted for degradation during senescence

The Sirtuin family of enzymes are key regulators of aging and senescence^4,5^. Previous studies show that several Sirtuin members are dynamically regulated during aging, however, these studies are primarily limited to SIRT1 and SIRT6, with limited study of SIRT7. We evaluated SIRT7 levels in three models of senescence (oncogene, etoposide, and replicative-induced) and observed via immunoblot analysis a global SIRT7 loss as cells undergo senescence (**Fig. 1A**, **Fig. 1B**, **Supp Fig. 1A**). While previous studies indicate that targeted depletion of SIRT7 can initiate senescence through ribosomal RNA (rRNA) stability^13^ and suppression of LINE-1 elements in human mesenchymal stem cells^15^, our observation that SIRT7 is naturally lost during senescence suggests a role in establishment of senescence.

**Figure 1:**
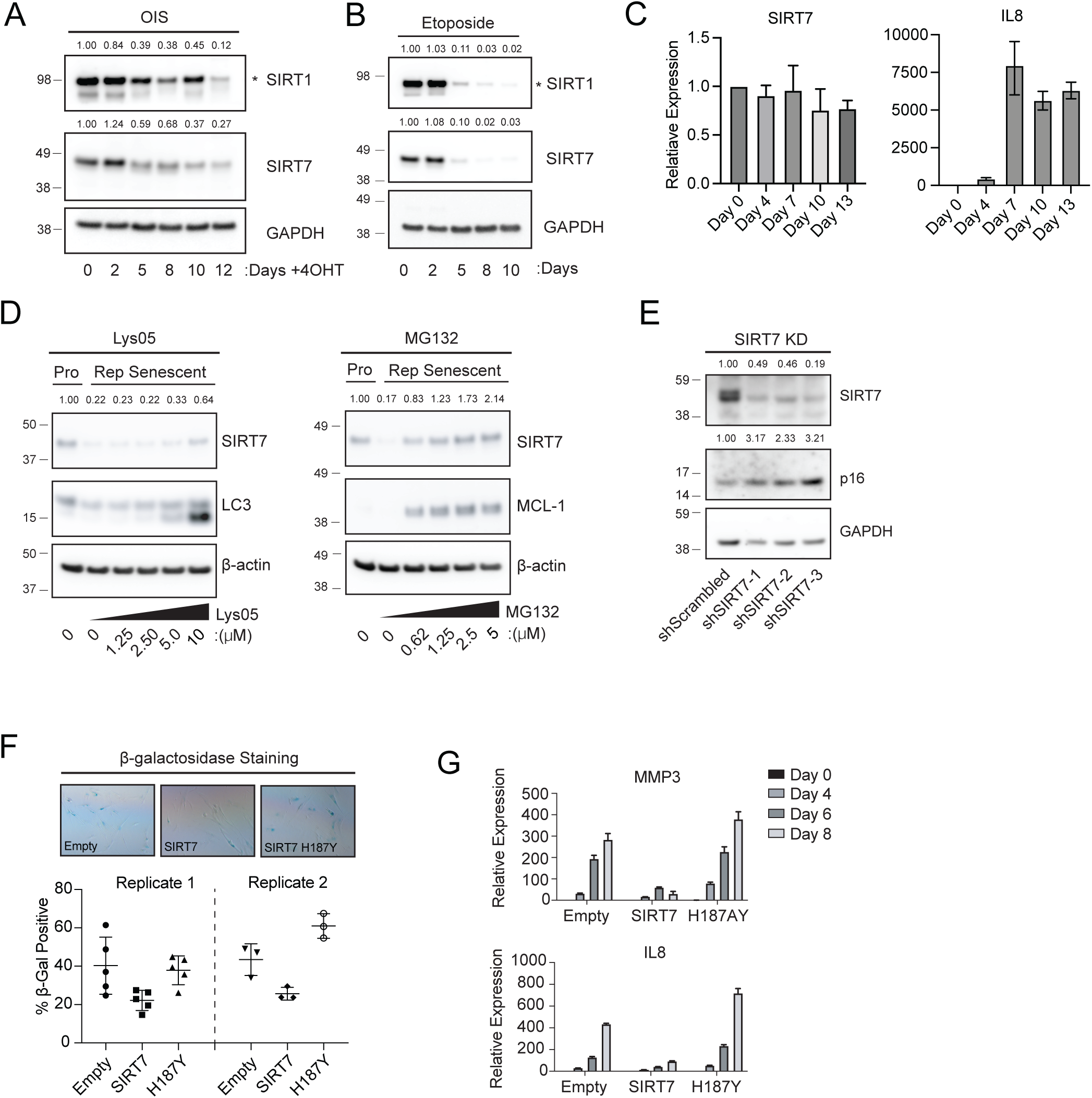
SIRT7 is targeted for degradation during senescence. A. Immunoblot of SIRT1 (targeted band is denoted by *) and SIRT7 during oncogene induced senescence (OIS). Cells were treated with the 4-Hydroxytamoxifen (4-OHT) to induce HRAS expression and harvested at multiple timepoints. GAPDH is used as a loading control. Intensity was quantified by adjusting to loading control (GAPDH) and normalized to Day 0. B. Immunoblot of SIRT1 (targeted band is denoted by *) and SIRT7 during etoposide induced senescence. Cells were treated with 100uM of etoposide for 48 hours and recovered in untreated media for an additional 8 days. GAPDH is used as a loading control. Intensity was quantified by adjusting to loading control (GAPDH) and normalized to Day 0. C. RT-PCR of SIRT7 (Left) and SASP gene (IL8) during etoposide induced senescence. Cells were treated with 100µM of etoposide for 48 hours and recovered in untreated media for an additional 8 days. Values represent standard deviation of two biological replicates and normalized to Day 0. D. Immunoblot of proliferative (Pro) or replicative senescent (Rep senescent) cells treated with increasing concentration of autophagy inhibitor Lyso5 (left) or proteasome inhibitor MG132 (right). LC3 and MCL-1 are used to quantify inhibition of autophagy or protease activity respectively. β-actin is used as a loading control. Intensity was quantified by adjusting to loading control (β-actin) and normalized to proliferative sample. E. Immunoblot of IMR90 cells depleted of SIRT7 via shRNA. GAPDH is used as a loading control. Intensity was quantified by adjusting to loading control (GAPDH) and normalized to scrambled control. F. Top – representative images of β-galactosidase staining of IMR90 cells overexpressing WT (SIRT7) or catalytically inactive (H187Y) SIRT7 on Day 13 of etoposide induced senescence. Bottom – Quantification of percent B-galactosidase for two independent replicates. Error bars represent standard deviation across technical replicates. G. RT-PCR of SASP genes MMP3 (Left) and IL8 (Right) of IMR90 cells overexpressing either empty, wildtype (SIRT7) or catalytically inactive (H187Y) SIRT7 during etoposide induced senescence. Cells were treated with etoposide for 24 hours and recovered for an additional 6 days. Cells were harvested at multiple timepoints and assessed for SASP genes.

Previous work demonstrates SIRT1 loss during senescence^8^ and consistently we found depletion of SIRT1 (**Fig. 1A**, **Fig. 1B**, **Supp Fig. 1A**). Senescence-mediated loss of SIRT1 is regulated post-translationally^8^, hence we assessed transcript levels of SIRT7 during etoposide-induced senescence (**Fig. 1C**) and examined published datasets in oncogene and replicative senescence (**Supp Fig. 1B**), and found constant SIRT7 transcript levels in proliferative and senescent cells, thus demonstrating post-translational regulation during senescence.

In our previous study, SIRT1 protein levels were reduced during senescence through nuclear-initiated autophagy^8^. To determine whether nuclear autophagy is relevant for SIRT7, we utilized a small molecule inhibitor of autophagy, Lys05^29^, but found little change in SIRT7 levels upon Lys05 treatment (**Fig. 1D left**, **Supp Fig. 1C**). These findings indicated a distinct mechanism of SIRT7 protein reduction during senescence. We tested the ubiquitination-mediated proteolysis pathway utilizing a proteasome inhibitor, MG132. SIRT7 protein levels showed rescue in senescent cells with increasing concentrations of MG132, reaching SIRT7 protein levels detected in proliferating cells (**Fig. 1D right**, **Supp Fig. 1C** and **1D**), and was independent of change in SIRT7 transcript levels upon MG132 inhibition (**Supp Fig. 1E**). Thus, SIRT7 may be targeted for proteasome degradation during senescence, and this is distinct from SIRT1 autophagic degradation.

Depletion of SIRT7 induces senescence^13,15^ and, consistently, we found that SIRT7 shRNA depletion in IMR90 cells led to increased expression of the key senescence marker p16 (**Fig. 1E**). We tested whether depletion of SIRT7 during senescence functions to, in particular, establish the senescence program, via expression of wildtype FLAG-SIRT7 (WT), catalytically inactive FLAG-SIRT7 (H187Y mutation^30^), or FLAG vector (control) in IMR90 fibroblasts undergoing etoposide-induced senescence. Upregulation of WT SIRT7 (but not catalytic inactive or vector) rescued senescence features of high β-galactosidase activity, high SASP expression (measured by MMP3 and IL8 expression) (**Fig. 1F**, **1G**). Additionally, we observed that upregulation of SIRT7 diminished other senescence phenotypes, such as reversing the loss of Lamin B1 in etoposide-induced senescence (**Supp Fig. 1F**) and lowering p16 and p21 expression in replicative senescence (**Supp Fig. 1G**). Hence, senescence suppression specifically requires SIRT7 and its deacetylation catalytic activity. Overall, these findings indicate that SIRT7 is selectively targeted by proteasome-mediated degradation during senescence, and this loss initiates the senescence program.

### SIRT7 loss during liver aging contributes to aging transcriptional changes

Accumulation of senescent cells contributes to tissue aging and selective targeting to eliminate these cells in mice promotes longevity and diminishes age-related disorders^31–35^. Because SIRT7 is reduced during cellular senescence (**Fig. 1**), we examined whether SIRT7 loss also occurs during tissue aging in mice. We quantified levels of SIRT7 in young and aged tissue samples from liver, kidney, muscle, lung, and heart. SIRT7 levels were lower in aged liver compared to young liver, whereas in contrast SIRT1 levels were constant (**Fig. 2A**, quantified **Fig. 2A right panels**). Further, SIRT7 levels were not decreased in other aged tissues (**Supp Fig. 2A-D**). We quantified RNA expression in young and old liver samples, and while senescence-associated p16 RNA was increased in aged livers, there were no significant changes in either SIRT1 or SIRT7 RNA levels (**Fig. 2B**). Together, these observations suggest that SIRT7 protein is reduced post-translationally during liver aging (unlike other tissues) and suggesting tissue-specific sirtuin protein regulation during aging.

**Figure 2:**
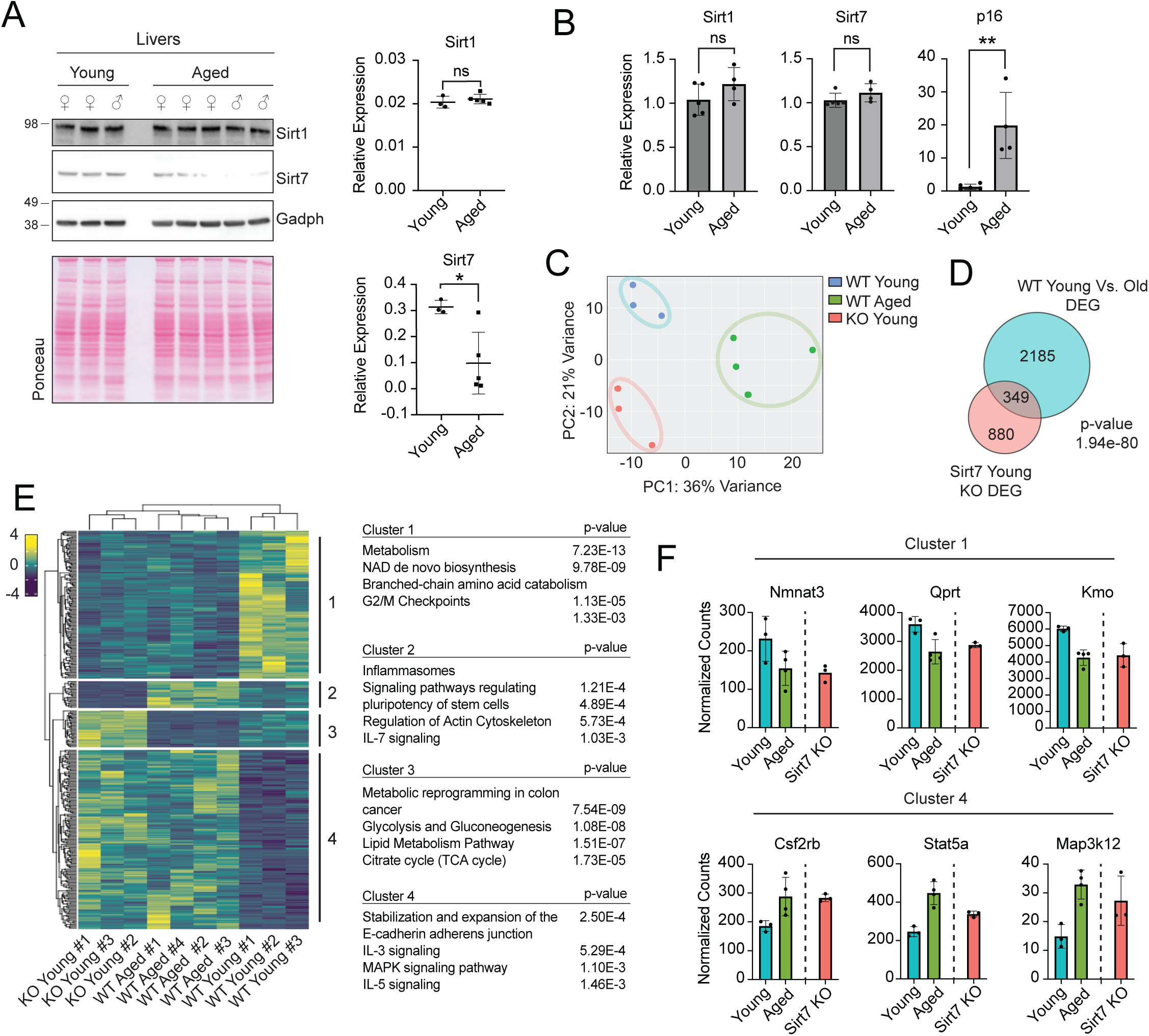
SIRT7 loss during liver aging contributes to aging transcriptional changes. A. Left - Immunoblot of SIRT1 and SIRT7 in young and aged mouse liver tissues. Ponceau is used as a loading control. Right – Quantification of band intensity normalized to GAPDH levels for SIRT1 and SIRT7. * p < 0.05 = unpaired t-test. B. RT-PCR of SIRT1 (left), SIR7 (middle) and p16 gene (right) in young and aged mice liver tissue. Samples were normalized to the median across young liver samples. Error bars represent standard deviation across liver samples. ** p < 0.01 = unpaired t-test. C. Principal component analysis (PCA) of wildtype (WT) young (blue), aged (green) and young SIRT7 KO (red) livers. D. Venn Diagram showing the overlap between WT aging DEGs (blue) and young SIRT7 KO DEGs (red). p-value calculated by hypergeometric probability. E. Left - Hierarchical clustering of overlapped genes (349 genes from Fig. 2D) of WT and SIRT7 liver samples with clusters labelled on the left. Values represent Z-score across liver samples. Right - Gene ontology (GO) for the clusters identified by hierarchical clustering. F. Normalized counts of Cluster 1 (top) or Cluster 4 (bottom) genes for WT young (blue), aged (green), and young SIRT7 KO (red) liver samples.

Previous studies in mice reveal that SIRT7 deletion accelerates aging^36^. To investigate how SIRT7 loss promotes aging, we examined transcriptome profiles of wildtype (WT) young, WT aged, and SIRT7 KO young mouse livers (**Supp Table 1**). In SIRT7 KO livers, there was no compensation from the other Sirtuin family members at the transcript level (**Supp Fig. 2E**). Further, in SIRT7 KO livers there was no global protein increase by western analysis of nuclear Sirtuin members SIRT1 or SIRT6 (**Supp Fig. 2F**). Broad assessment via principle component analysis (PCA) of the transcriptomics of SIRT7 WT young and old livers indicated distinct transcriptional profiles (**Fig. 2C** and **Supp Fig. 2G**): young livers had upregulated genes associated with metabolic pathways including amino acid and TCA cycles (**Supp Fig. 2H top**), whereas aged livers were upregulated for genes associated with immune response (**Supp Fig. 2H bottom**). As liver is a complex metabolic organ, metabolic changes accompanying aging may indicate disrupted energy metabolism and may contribute to liver-specific age-related disorders like insulin resistance. Additionally, the increase in immune associated genes in aged livers may relate to an increase of senescent hepatocytes which release inflammatory cytokines leading to inflammation and tissue dysfunction^37,38^.

To identify how SIRT7 contributes to aging, we compared young SIRT7 WT and KO liver transcriptomes by PCA, revealing that the young SIRT7 KO clustered distinctly from WT young and aged samples (**Fig. 2C**) with approximately 1229 differentially regulated genes (DEGs) between SIRT7 WT and KO young livers (**Supp Fig. 2I**). SIRT7 deacetylates histones, specifically modifications H3K18ac^30^ and H3K36ac^39^ leading to gene repression. Consistently we observed a slight preference for gene upregulation (688 genes) compared to downregulated (541 genes) in SIRT7 KO livers (**Supp Fig. 2I**). Genes that were downregulated in SIRT7 KO livers were associated with metabolism such as fatty acid regulation, glycolysis, and gluconeogenesis pathways, whereas genes that were upregulated in SIRT7 KO livers were associated with amino acid metabolism and NAD biosynthesis (**Supp Fig. 2J**). These observations are consistent with previous findings in SIRT7 KO mice of hepatic SIRT7 regulating lipid metabolism^11,40^, and our data suggest there may be additional metabolic reprogramming occurring with a shift away from glucose metabolism. As the liver is a pivotal organ in regulating metabolic homeostasis within the body^41^, these observations indicate SIRT7 is vital in maintaining appropriate metabolic transcriptional programs.

We searched for aging-associated SIRT7 regulated genes via overlapping genes that were differentially expressed in SIRT7 KO livers with genes dysregulated in liver aging. This revealed 349 intersecting genes, and these represented one-third of total SIRT7 KO DEGs (**Fig. 2D**). The expression of the overlapping DEGs across WT young, WT aged, SIRT7 KO young generated four unique clusters of expression patterns (**Fig. 2E**). Revealingly, hierarchical clustering showed that the SIRT7 KO young livers clustered with WT old livers, and there was distinct clustering of WT young livers. Clusters 1 and 4 showed highest similarity and the largest gene number between SIRT7 KO and WT aged; these were associated with metabolism, IL-3/5, and MAPK signaling (**Fig. 2E** and **Fig. 2F**, example of *Stat5a* gene in **Supp Fig. 2K**). Taken together, our data suggest that the rapid aging observed in the SIRT7 KO mouse could be due to loss of SIRT7’s ability to regulate expression of key metabolic, signaling and inflammatory pathways to protect against senescence and liver aging.

### SIRT7 interacts with epigenetic regulators and transcription factors

SIRT7 occurs in various cellular compartments with numerous potential protein interactions^6,42,43^, thus we explored SIRT7 interactors and their contribution to senescence upon reduction of SIRT7. We derived IMR90 fibroblast lines expressing FLAG tagged-SIRT7 (or FLAG alone) and performed immunoprecipitation (IP) for the FLAG epitope in conjunction with mass spectrometry (MS) in proliferating cells (schematic **Supp Fig. 3A**, **Supp Table 2**). Binding partners of SIRT7 were defined as proteins: (1) exclusively enriched in SIRT7 IP, (2) enriched two-fold over FLAG alone control, and (3) not identified as common contaminants in MS^44^ (**Supp Fig. 3A**). SIRT7 was the most significantly enriched protein, thus confirming efficient and specific SIRT7 IP **(Supp Fig. 3B** and **Supp Fig. 3C**). Overall, there were 533 SIRT7-interacting partners, and, via cross-comparison with published SIRT7 interactome datasets^15,45,46^, we identified a cohort of shared proteins (**Fig. 3A**). We noted 323 proteins exclusive to our study, and since SIRT7 was exogenously expressed in HEK293T cells in previous datasets, these unique proteins may be cell type specific SIRT7 interactors^15,45,46^. There was a cohort of proteins shared across the datasets, including the E3 Ubiquitin Ligase, TRIP12, and ribosomal processing proteins FBL and TEX10 (**Fig. 3A**).

**Figure 3:**
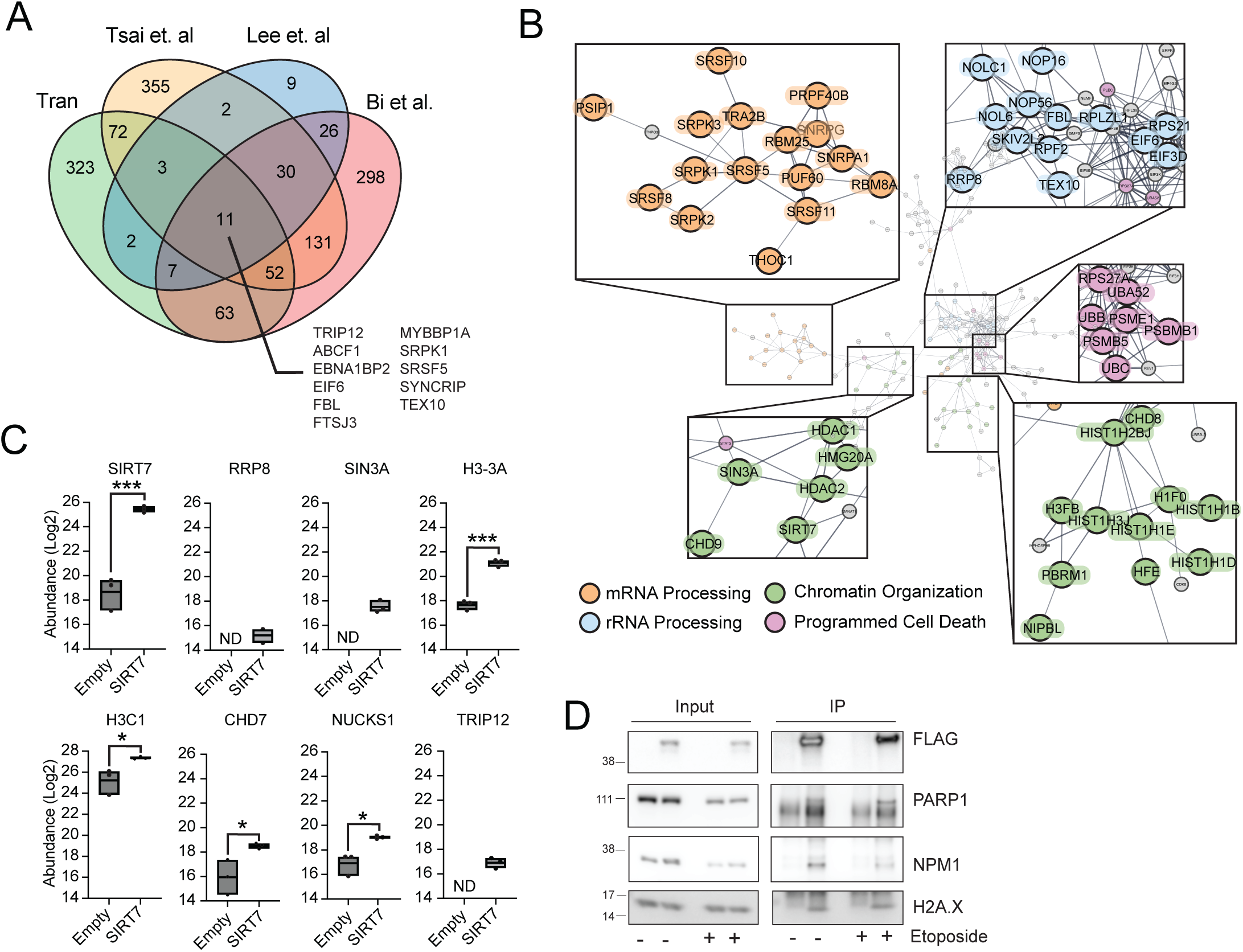
SIRT7 interacts with epigenetic regulators and transcription factors. A. Venn diagram of immunoprecipitation of SIRT7 in IMR90 (Tran) compared to published SIRT7 IP in 293T. Overlap SIRT7 interacting proteins between studies are listed. B. SIRT7 interaction network generated by STRING-DB using filtered SIRT7 interactors. Zoomed in image of selected subnetworks associated with mRNA processing (orange), ribosomal processing (blue), chromatin organization (green), and cell death (pink). Darkness of edges corresponds to the confidence of interaction. C. Box plots of peptide abundance of selected proteins enriched in either FLAG-Empty or FLAG-SIRT7 immunoprecipitation. Interior line within box plot corresponds to mean across replicates. *** p < 0.001 * p < 0.05 = unpaired t-test. D. Immunoprecipitation of FLAG in IMR90 cells expressing either FLAG-Empty or FLAG-SIRT7 in proliferating or etoposide-induced senescent cells and immunoblotted for components of H2AX complex. ND denotes not detected by mass spectometry.

We employed STRING-DB network analysis^47^ to identify putative SIRT7 complexes and functional SIRT7 interactors. We identified several nucleolar interactors involved in ribosomal RNA (rRNA) processing, as previously reported for SIRT7^48,49^ (**Fig. 3B blue**), and proteins in mRNA processing (such as SRSF family) (**Fig. 3B orange**) suggesting a SIRT7 role in mRNA splicing and processing. Other interactors were chromatin associated proteins (**Fig. 3B green**), underscoring SIRT7’s gene regulation role, including several histone deacetylases (HDAC1, HDAC2, and SIN3A), suggesting coordinated chromatin compaction with SIRT7 (**Fig. 3B green** and **Fig. 3C**). We utilized gene enrichment analysis and identified additional pathways of SIRT7’s binding partners including STAT3 and WNT signaling, epigenetic regulation of rRNA and nicotinamide (NAD+) metabolism (**Supp Fig. 3D**). Indeed, SIRT7 is a NAD+ dependent enzyme, and thus binding of numerous NAD enzymes (e.g. NNMT, NMNAT1, and NAMPT) suggests unique metabolic-epigenetic crosstalk to supply cofactors for SIRT7’s catalytic function (**Supp Fig. 3D** and **Supp Fig. 3E**). Several chromatin complexes were associated with SIRT7, including the SIN3A complex and the H2AX complex (**Fig. 3C** and **Supp Fig. 3E**) and these were confirmed by FLAG-SIRT7 IP-western (**Fig. 3D**). Taken together, SIRT7 interacts with a wealth of metabolic and chromatin-relevant complexes and pathway and these interactions may be pivotal in the role of SIRT7 in the establishment of senescence.

### SIRT7 deacetylation of NUCKS1 promotes chromatin binding

SIRT7 deacetylates histone substrates H3K18ac and H3K36ac to elicit gene repression^30,39^. We noted proteomic interactions of SIRT7 with transcription factors. One of these was NUCKS1 (nuclear casein kinase and cyclin-dependent kinase substrate 1) (**Fig. 3C bottom**), a predicted high mobility group (HMG) protein associated with gene activation and cell proliferation, DNA damage, NF-kB activation, and metabolism^25–28,50,51^. Given the overlap of these processes with senescence, we investigated the SIRT7-NUCKS1 relationship during cellular senescence.

We performed IP of endogenous SIRT7 and detected NUCKS1 (**Fig. 4A left**, NUCKS1 antibody validated in **Supp Fig. 4A**), and the reverse IP of endogenous NUCKS1 detected SIRT7 by immunoblot (**Fig. 4A right**). NUCKS1 levels are dynamically regulated during cell cycle and upon DNA damage^25,27,28^, however, we found no change of NUCKS1 RNA (**Supp Fig. 4C**) or protein (**Supp Fig. 4B**) in etoposide-induced senescence compared to proliferation, or in public transcriptome datasets of oncogene- and replicative-induced senescence (**Supp Fig. 4D**). NUCKS1 is dynamic in the nucleus and onto chromatin in cell cycle^25^ or during DNA repair^27^, thus, we evaluated NUCKS1 localization during cellular senescence and found increased NUCKS1 colocalization with nuclear maker DAPI in senescent cells by immunofluorescence (**Fig. 4B**) and confirmed using a second NUCKS1 antibody (**Supp Fig. 4E**). We examined chromatin binding via cellular fractionation/compartmental proteomics-mass spectrometry of proliferating and etoposide induced senescent IMR90 cells (**Fig. 4C left**). NUCKS1 levels were higher in the chromatin fraction in senescent cells compared to proliferating, without changes of NUCKS1 levels in the cytoplasmic or nuclear fractions (**Fig. 4C right**), and higher chromatin levels of NUCKS1 in senescence was confirmed by immunoblot analysis (**Fig. 4D**). Overall, these findings indicate increased chromatin association of NUCKS1 during senescence.

**Figure 4:**
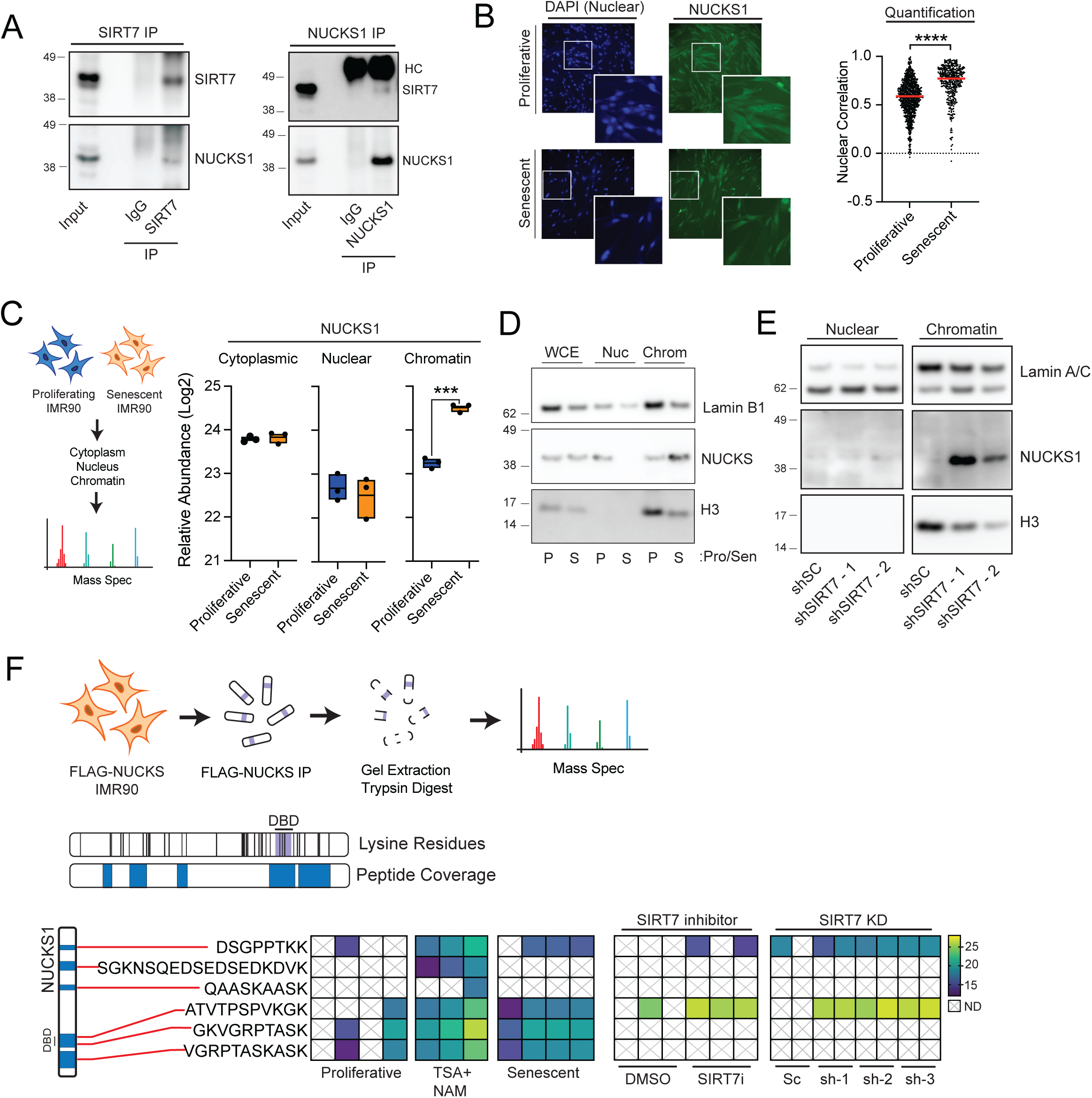
SIRT7 deacetylation of NUCKS1 promotes chromatin binding. A. Immunoprecipitation of either IgG, SIRT7 (left) or NUCKS1 (right) in proliferation IMR90 and immunoblotted for either SIRT7 or NUCKS1. B. Left - Representative immunofluorescence images of NUCKS1 (green) localization in proliferating (top) and senescent (bottom) cells. DAPI (blue) is used to show nuclear staining. Right - Quantification of DAPI nuclear staining with NUCKS1 signal from immunofluorescence images. Each dot represents an individual nuclei with red horizontal bars indicating median value across all nuclei. **** p < 0.0001 = unpaired t-test. C. Left - Schematic of compartmental proteomics in proliferating and senescent cells. Right - Enrichment of NUCKS1 peptides in different subcellular compartments in proliferating (blue) and etoposide induced senescent cells (orange). *** p < 0.0001 = unpaired t-test. D. Immunoblot of NUCKS1 in either whole cell extract (WCE), nuclear (Nuc), and Chromatin (Chrom) fractions in proliferative or etoposide induced senescence IMR90 cells. Lamin B1 depletion is used as a marker of senescence and H3 is used as a control for nuclear versus chromatin fractionation. E. Immunoblot of NUCKS1 in either the nuclear (left) or chromatin (right) fraction in control (shSC) or shRNA depleted SIRT7 IMR90 cells (shSIRT7 - 1/2). Lamin A/C and H3 are used as controls for nuclear and chromatin fractionation. F. Top - Schematic of acetylated residues of NUCKS1 by mass spectrometry. FLAG-NUCKS1 was immunoprecipitated in IMR90 cells and was gel purified followed by trypsin digestion before mass spectrometry identification. Middle - Representation of NUCKS1 protein with lysine residues denoted by black vertical bars (top) and DNA binding domain (DBD) highlighted in purple. Blue boxes (bottom) represent location of peptides detected in mass spectrometry analysis. Bottom - Heatmap showing the peptide abundance of acetylated peptides identified from mass spectrometry in DMSO (Proliferating), TSA+NAM, etoposide-induced senescent cells, or SIRT7 manipulation. Each column corresponds to an independent replicate. White boxes with “X” indicate peptides not identified by the mass spectrometry. Relative location of acetylated NUCKS1 peptide is shown by the red line matching the peptide with the NUCKS1 protein (left). Line corresponds to the putative DNA binding domain.

The dual findings of SIRT7 interaction with NUCKS1 and the recruitment of NUCKS1 onto chromatin during senescence led us to hypothesize that the targeted proteasome degradation of SIRT7 during senescence leads to NUCKS1 recruitment onto chromatin. To test this, we depleted SIRT7 using shRNA in IMR90 cells followed by fractionation to examine NUCKS1 localization. While NUCKS1 transcript and protein levels remained unchanged (**Supp Fig. 4F** and **Supp Fig. 4G**), SIRT7 depletion led to higher levels of NUCKS1 within the chromatin fraction (**Fig. 4E**). Orthogonally, we treated IMR90 cells with a SIRT7 inhibitor^52^ which led to an increase in SIRT7 deacetylation target H3K36ac levels in a dose-dependent manner (but did not change SIRT7 levels) (**Supp Fig. 4H**), and concomitantly increased NUCKS1 specifically within the chromatin fraction without overall change in levels (**Supp Fig. 4I**). In sum, this suggests SIRT7 loss can promote NUCKS1 localization onto chromatin.

NUCKS1 protein is heavily post-translationally modified (PTM) by acetylation, phosphorylation, methylation, and formylation^53^. In particular, NUCKS1 is highly enriched for lysine residues with a large number near to, and within the DNA binding domain (**Fig. 4F**, middle). NUCKS1 is phosphorylated by CK2^54^, however NUCKS1 acetylation dynamics and function are under studied. To explore this, we expressed FLAG-NUCKS1 in IMR90 cells and immunoprecipitated for FLAG for proteomic analysis to identify NUCKS1 PTMs (**Fig. 4F top**). To identify putative acetylated residues, we treated cells with deacetylase inhibitors [Nicotinamide (NAM) and Trichostatin A (TSA)] to enrich for NUCKS1 acetylation. Proteomics of NUCKS1 had ∼30% peptide coverage with the best representation in the vicinity of the DNA binding domain. We identified 5 peptides with acetylated lysines (**Fig. 4F middle**, **Supp Table 3**), and evaluated these in proliferating and senescent cells, finding increased senescence-associated acetylated peptides surrounding the DNA binding domain (**Fig. 4F bottom left**). To examine the consequence of increased NUCKS1 acetylation on cellular distribution, we inhibited protein deacetylase using a combination of TSA+NAM in proliferating IMR90 cells and while we found little to no difference in NUCKS1 levels upon treatment (**Supp Fig. 4J**), we found increased nuclear localization (**Supp Fig. 4K**). Furthermore, deacetylase inhibition increased levels within the chromatin fraction (**Supp Fig. 4L**), hence NUCKS1 acetylation increased NUCKS1 chromatin binding.

The findings relating NUCKS1 acetylation and chromatin binding led us to hypothesize that SIRT7 deacetylates NUCKS1. To test this, we enriched for FLAG-NUCKS in proliferating IMR90 cells that were treated with SIRT7 inhibitor or depleted of SIRT7 by shRNA. Interestingly, we observed acetylation of peptides in the N-terminal and DNA binding domain of NUCKS1 upon SIRT7 inhibition or KD, and these were differentially acetylated between proliferative and senescent cells (**Fig. 4F bottom right**). Collectively, these findings show chromatin recruitment of acetylated NUCKS1 during senescence, mediated by SIRT7 proteolysis.

### HMG protein NUCKS1 regulates senescence

NUCKS1 is a transcription factor involved in cell cycle and inflammatory response, hence we hypothesized that SIRT7 regulation of senescence could be mediated through NUCKS1. We assessed etoposide-associated senescence following depletion of NUCKS1 in IMR90 cells using three independent shRNAs (**Supp Fig. 4A**), finding decreased β-galactosidase staining cells (**Fig. 5A**) and reduced expression of SASP genes, IL8 and MMP3 (**Fig. 5B**). Conversely, overexpression of FLAG-NUCKS1 (**Supp Fig. 5A**) exacerbated etoposide-associated senescence in increased β-galactosidase cells and higher SASP gene expression (**Supp Fig. 5B** and **Supp Fig. 5C**).

**Figure 5:**
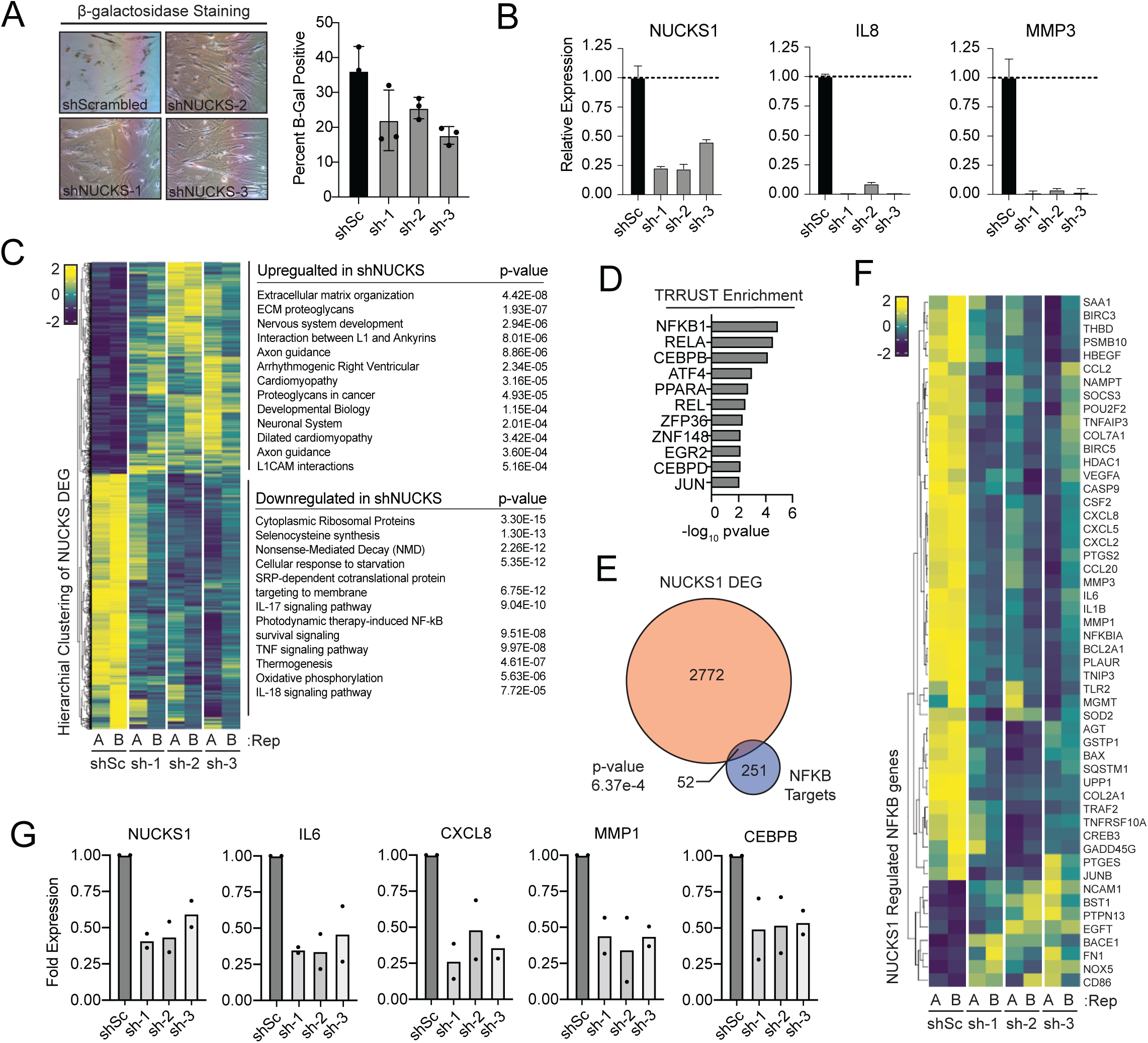
NUCKS1 involved in NFKB and CEBPB transcriptional programs. A. Left – representative images of b-galactosidase staining of either WT (shScrambled) or NUCKS1 depleted (sh-1/2/3) IMR90 cells on day 13 of etoposide induced senescence. Right - Quantification of percent B-galactosidase. Error bars represent standard deviation across three biological replicates. B. RT-PCR of NUCKS1 and SASP gene (IL8 and MMP3) of either WT or NUCKS1 depleted IMR90 cells on day 13 of etoposide induced senescence. Values represent standard deviation of three technical replicates and normalized to scrambled control. C. Left - Heatmap of DEGs between shScrambled and shNUCKS in etoposide induced senescent IMR90 cells (2738 genes). Values represent Z-score across each replicate. Right - Gene ontology (GO) for either upregulated (top) or downregulated (bottom) in NUCKS1 depleted cells. D. TTRUST (Transcriptional Regulatory Relationships Unraveled by Sentence-based Text mining) enrichment of NUCKS1 DEGs. E. Venn Diagram showing the overlap between NUCKS1 DEGs (orange) and NFKB targets from TRRUST database (blue). F. Heatmap of overlapped genes (52 genes from Fig. 5E) of NUCKS1 DEG and NFKB targets. Values represent Z-score across each replicate shScrambled or shNUCKS1 depleted cells. G. Expression of NFKB and CEBPB targets regulated by NUCKS1 in etoposide induced senescent IMR90 cells. Values represent fold difference to shScrambeld control.

Given the role of NUCKS1 in transcription, we performed transcriptome analysis of NUCKS1 KD cells in etoposide-associated senescence and identified ∼7-fold more differentially expressed genes (DEGs) in senescent compared to proliferative cells (2824 vs 402 DEGs) with few overlapping DEGs (**Supp Fig. 5D**, **Supp Table 4**). This indicates that NUCKS1 has a regulatory role in senescence, coinciding with increased binding of NUCKS1 onto chromatin in senescent cells (**Fig. 4D** and **Fig. 4F**). NUCKS1 KD in proliferative cells led to more downregulated than upregulated genes (255 vs 147) and DEGs were associated with ECM remodeling, Vitamin D signaling and adipogenesis (**Supp Fig. 5E** and **Supp Fig. 5F**). Examination of senescent associated NUCKS1 KD DEGs indicate roughly equivalent upregulated and downregulated genes (**Fig. 5C**, **Supp Fig. 5G**); the upregulated genes were associated with extracellular matrix and neuronal pathways, while downregulated genes (genes that are upregulated by NUCKS1 in senescence) were involved in ribosomal pathways, inflammatory signaling and stress response (**Fig. 5C right**). Interestingly, these latter processes are linked to the senescence program and thus suggest that NUCKS1 mediates transcriptional upregulation of certain senescence-pathways.

As transcription factors often work in concert, we explored whether other transcription factors contribute to NUCKS1 mediated gene regulation, via TRRUST enrichment for motif analysis^55^. This pointed to significant enrichment of NFkB and CEBPβ motifs specifically within senescence-associated genes altered in NUCKS1 KD (**Fig. 5D**, **Fig. 5E**, and **Supp Fig. 5H**). Interestingly, NFkB and CEBPβ are well-known regulators of senescence and specifically upregulate SASP gene expression^56–58^. Indeed, we focused on NFkB target genes within the NUCKS1 KD DEGs (52 gene overlap in **Fig. 5E**) and the majority of these were downregulated upon depletion of NUCKS1 in senescence (**Fig. 5F**). Similarly, CEBPβ target genes that overlapped NUCKS1 DEGs (18 gene overlap in **Supp Fig. 5H**) were predominantly downregulated upon NUCKS1 loss (**Supp Fig. 5I**). Further, CEBPβ itself decreased expression upon NUCKS1 KD in senescent cells (**Fig. 5G**) which could explain diminished expression of CEBPβ target genes. Unlike CEBPβ, transcript levels of NFKB1 or RELA were not lowered upon NUCKS1 KD in senescent cells (**Supp Fig. 5J**), which suggests a direct role of NUCKS1 in regulating NFkB signaling or targeting of NFkB to specific genes. In sum, these observations indicate that NUCKS1 is a regulator of senescence and contributes to the senescence transcriptional program.

### SIRT7 KO livers have elevated chromatin bound NUCKS1

We showed above that SIRT7 protein levels are reduced in the livers of aged mice (see **Fig. 2**), and that SIRT7 loss in senescent IMR90 cells leads to increased NUCKS1 chromatin association (see **Fig. 4E**). We next examined NUCKS1 transcriptionally and at the protein level, between young and aged livers from SIRT7 WT and knockout (KO) mice. As expected, Sirt7 transcripts were undetectable in the knockout livers, however, Nucks1 RNA levels were unchanged (**Fig. 6A**). We examined NUCKS1 protein levels, first validating antibody specificity in NUCKS1 depleted mouse T lymphoblast cells (EL4 cells) (**Supp Fig. 6A** and **Supp Fig. 6B**). Interestingly, NUCKS1 protein was increased in the SIRT7 KO livers compared to WT livers derived from both young and aged mice (**Fig. 6B**, **Supp Fig. 6C**). These observations suggest that in mouse liver SIRT7 may regulate NUCKS1 post-translationally.

**Figure 6:**
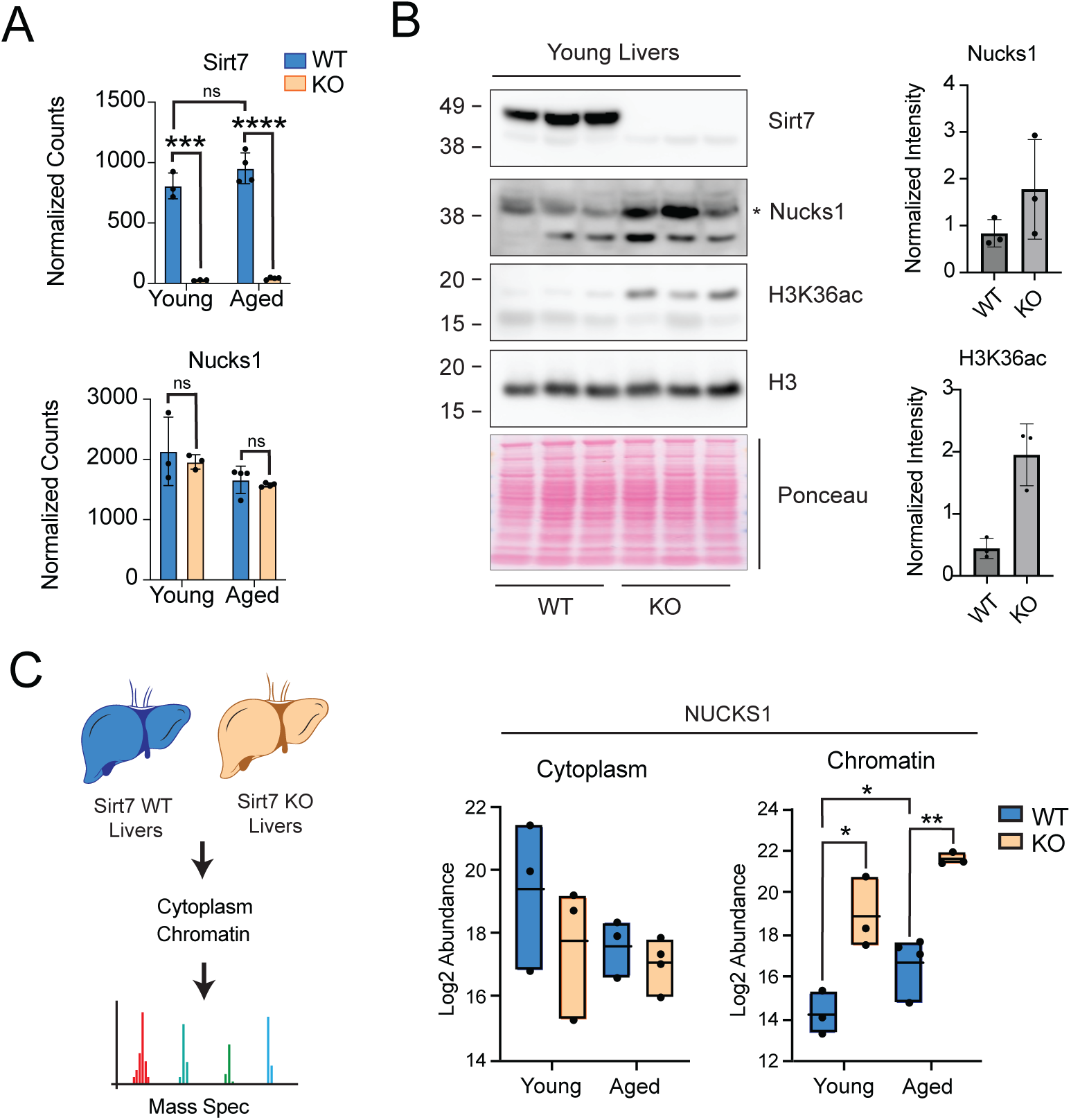
SIRT7 KO livers have elevated chromatin bound NUCKS1. A. Normalized counts for Sirt7 and Nucks1 in young and ages WT and Sirt7 KO livers. **** p < 0.0001 *** p <0.001 = unpaired t-test. B. Left - Immunoblot of WT or KO Sirt7 in young livers (Nucks1 targeted band is denoted by *). Ponceau staining is used as a loading control. Right - Quantification of Nucks1(Top) and H3K36ac (Bottom) in WT and KO Sirt7 young livers. Intensity was quantified by adjusting to H3 and normalized to median across all samples. C. Left - Schematic of compartmental proteomics in young and aged livers. Right - Enrichment of Nucks1 peptides in different subcellular compartments in WT (Blue) or KO (Orange) Sirt7 young or aged livers. ** p < 0.01 * p < 0.05 = unpaired t-test.

We showed that NUCKS1 is bound to chromatin during senescence where loss of SIRT7 promotes NUCKS1 chromatin recruitment (see **Fig. 4D**, **4E**, and **4F**). Hence, the elevated levels of NUCKS1 in SIRT7 KO livers may also manifest within chromatin. To determine whether NUCKS1 is redistributed during liver aging, we performed compartmental fractionation of either WT or KO SIRT7 in young and aged mice in conjunction with mass spectrometry (**Fig. 6C left** and **Supp Fig. 6D**). NUCKS1 levels in the cytoplasm trended lower during aging both in the WT and SIRT7 KO mice (**Fig. 6C right**); in contrast NUCKS1 levels in the chromatin fraction showed significant increase in aging and most strongly in the KO livers (**Fig. 6C right**). We particularly note the strong elevation from young to old livers in the SIRT7 KO mouse (**Fig. 6C right**). These observations indicate that NUCKS1 increases within the chromatin fraction during liver aging and this shift in localization is promoted by loss of SIRT7.

### NUCKS1 genome-binding corresponds with liver aging associated genes

We observed NUCKS1 increased binding to chromatin in aged mouse livers, exacerbated by *Sirt7* deletion (see **Fig. 6C right**). Thus, we investigated how NUCKS1 binding impacts gene expression during aging and SIRT7 loss, utilizing published ChIP sequencing dataset of NUCKS1 binding in isolated young hepatocytes compared to activation-associated histone modifications at these sites in the liver^50,59,60^. Consistent with previous studies^50^, our analysis showed NUCKS1 binding peaks at active promoters associated with H3K4me3 and activation modifications H3K9ac and H3K27ac (**Fig. 7A** cluster 1 (top); **Supp Fig. 7A**). For example, binding of NUCKS1 and histone modifications occurs at the promoter of *Stat5a* (**Fig. 7B lower blue bar**). We also found NUCKS1 peaks associated with active enhancers coinciding with H3K4me1 and H3K27ac peaks (**Fig. 7A** cluster 2 (bottom); **Supp Fig. 7A**); this binding occurs at the enhancers of Stat5a (**Fig. 7B lower orange bars**).

**Figure 7:**
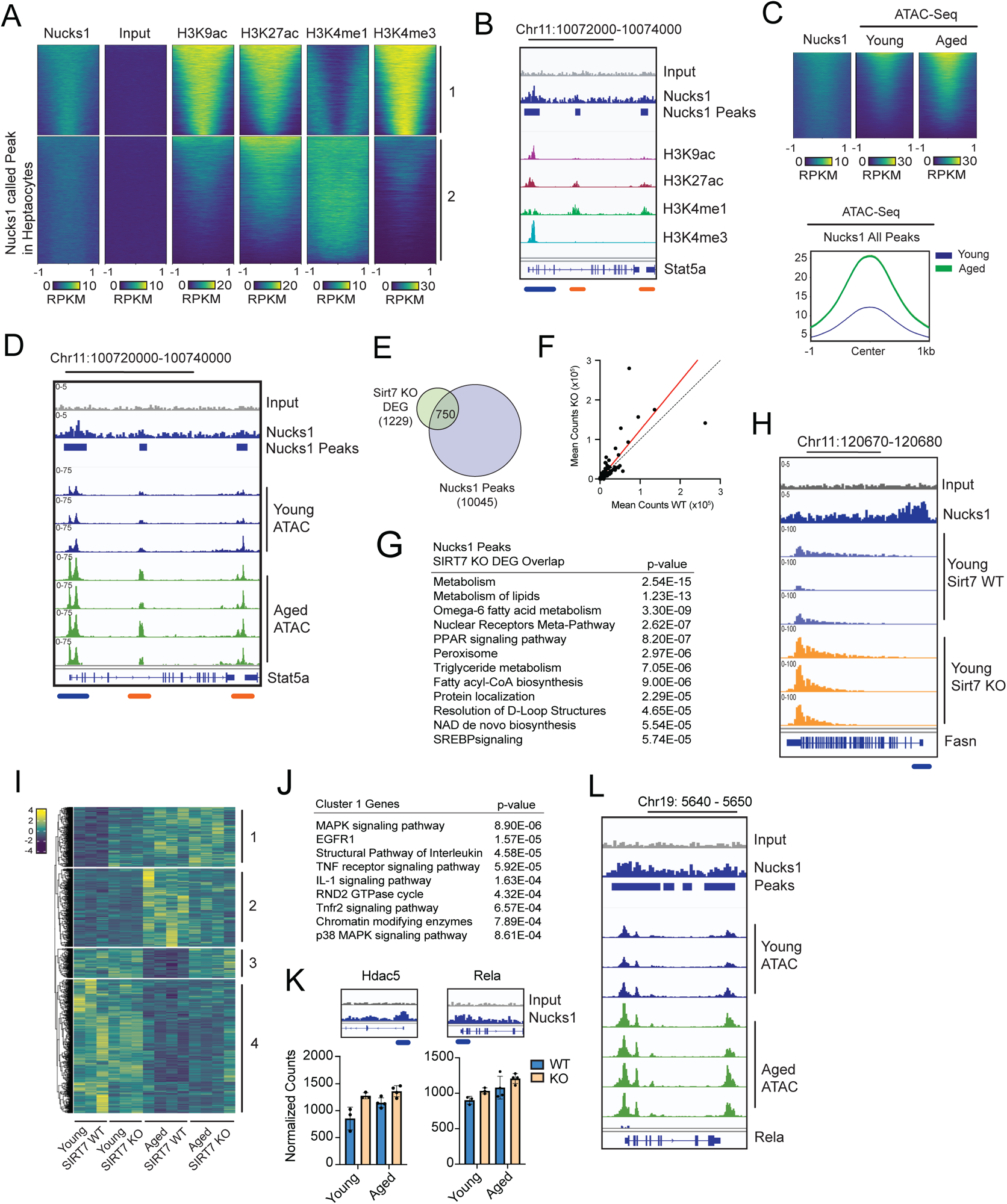
NUCKS1 genome-binding corresponds with liver aging associated genes. A. Heatmap of ChIP signal for Nucks1 called peaks for Nucks1, input (control), H3K9ac, H3K27ac, H3K4me1, and H3K4me3 in liver tissue. Peaks were k-means clustered into two distinct clusters. B. Gene tracks of ChIP-Seq signal for input (grey), Nucks1 (blue), Nucks1 called peaks (blue bars), H3K9ac (purple), H3K27ac (maroon), H3K4me1 (green), and H3K4me3 (blue) at the Stat5a locus. Blue lines underneath tracks indicate promoter region while orange lines indicate putative enhancers. C. Top - Heatmap of ChIP signal for Nucks1 called peaks for Nucks1, input and ATAC-Seq in young and aged livers. Metaplot of ATAC-Seq signal at Nucks1 called peaks in young (blue) and aged (light green) livers. D. Gene tracks of ChIP-Seq signal for input (grey), Nucks1 (blue), Nucks1 called peaks (blue bars), and ATAC-Seq signal in young and aged livers at the Stat5a locus. Blue lines underneath tracks indicate promoter region while orange lines indicate putative enhancers. E. Venn diagram of Nucks1 called peaks and differentially expressed genes (DEGs) in Sirt7 WT and KO young livers. F. Expression plot of normalized counts for Sirt7 WT (x-axis) and KO (y-axis) livers for overlapped genes (750 genes from Fig 7D). Linear regression of data points is denoted by red line. Black dotted lines represent regression of 1. G. Gene ontology for overlapped genes (750 genes from Fig 7D) between Nucks1 peaks in hepatocytes and DEGs between Sirt7 WT and KO livers. H. Gene tracks of ChIP-Seq signal for input (grey), Nucks1 (blue) and RNA expression in young Sirt7 WT (light blue) and Sirt7 KO (orange) livers at the Fasn locus. Blue lines underneath tracks indicate promoter region. I. Hierarchical clustering of overlapped genes (1778 genes from Fig 7E) of Nucks1 peaks and DEGs between young and aged livers for expression in Sirt7 WT or KO young and aged livers. Clusters are denoted on the left side of the heat map. J. Gene ontology for cluster 1 genes defined in Fig 7H. K. Top - Gene tracks of ChIP-Seq signal for input (grey) and Nucks1 peaks (blue) for target genes. Bottom - Expression of target genes in Sirt7 WT and KO in young and aged livers. L. Gene tracks of ChIP-Seq signal for input (grey), Nucks1 (blue), Nucks1 called peaks (blue bars), and ATAC-Seq signal in young and aged livers at the Rela locus.

Thus, NUCKS1 is associated with gene promoters and enhancers in the young basal state, and, based on our findings above in senescence and aging (**Fig. 4E** and **Fig. 6C**), we predict increased NUCKS1 ChIP binding and enhanced transcription activity during liver aging. To address this, we examined chromatin changes around NUCKS1 bound sites during aging, utilizing published ATAC-seq in aged mouse livers^61^. We found that a large proportion of NUCKS1 peaks (from young basal state; **Fig. 7A**) had increased chromatin accessibility in aged livers compared to young (**Fig. 7C**, **Fig. 7D**, and **Supp Fig. 7B**); increased NUCKS1 binding to chromatin observed during aging could lead to this increased accessibility. Interestingly, we observed increased accessibility in aged livers at NUCKS1 bound sites (**Supp Fig. 7B**).

Our previous observations indicate NUCKS1 chromatin localization was increased upon SIRT7 KD and in the livers of SIRT7 KO mice (**Fig. 4F** and **Fig. 6C**), suggesting that a subset of the NUCKS1 peaks (**Fig. 7A**) may show increased binding and thus higher gene expression in SIRT7 KO livers. We examined NUCKS1 peaks (**Fig. 7A**) associated with SIRT7 DEGs in young livers and identified a cohort of 750 genes with NUCKS1 binding and expression changes upon *Sirt7* deletion (**Fig. 7E**). These genes tend to be more highly expressed upon Sirt7 deletion (**Fig. 7F**, **Supp Fig. 7C**) and are associated with metabolic processes such as lipid metabolism (**Fig. 7G**) and include genes like *Fasn*, Fatty acid synthase (**Fig. 7H**). Thus, SIRT7 regulation of hepatic lipid metabolism^11^ may involve NUCKS1 binding to mediate expression changes of these metabolic genes.

We correlated NUCKS1 binding with aging-associated genes via overlap of NUCKS1 peaks in young hepatocytes with DEGs in liver aging, revealing ∼1800 age-associated DEGs with a NUCKS1 peak either at the promoter or in gene proximity (**Supp Fig. 7D**). Hierarchical clustering across SIRT7 WT and KO in young compared to aged livers indicated 4 distinct DEG clusters, with the majority of genes defined by age (**Fig. 7I**, clusters 2 and 4). Interestingly, cluster 1 contained a subset of upregulated genes in liver aging that were also elevated in the SIRT7 KO young (**Fig. 7I**, **Supp Fig. 7E**); these could represent a gene class with elevated NUCKS1 binding upon *Sirt7* deletion and could be key to rapid aging in the SIRT7 KO mouse. Indeed, GO analysis of cluster 1 reveals MAPK signaling, inflammatory response, and chromatin remodeling (**Fig. 7J**). Chromatin regulators within cluster 1 included *Hdac5* and *Dot1l*, which could contribute to gene expression changes during liver aging (**Fig. 7K**, **Supp Fig. 7F**). Of particular interest was the cohort of MAPK-related genes (**Supp Fig. 7F**), as MAPK pathways are linked to senescence phenotypes, including NFKB-mediated SASP expression and epigenetic regulation^62–65^. Interestingly, genes in MAPK signaling included several MAP kinases and *Rela* (**Fig. 7K**, **Supp Fig. 7F**). *Rela* and *Nfkb1* are key regulators of SASP expression^56,58^ and we found NFKB targeted genes strongly associated with NUCKS1 depletion in senescent IMR90s (see **Fig. 5D**). To further explore how Nucks1 bound sites were altered in liver aging, we utilized the ATAC-seq data from young and aged livers^61^ revealing increased accessibility of the *Rela* locus during aging (**Fig. 7L**).

Overall, these observations indicate that Nucks1 binds near enhancers and promoters of genes that tend to have increased accessibility during liver aging. Further, NUCKS1 peaks occur near a subset of aged-associated genes in the liver that are regulated by SIRT7, including the key SASP regulator *Rela*.

## DISCUSSION

In this study, we show that SIRT7 is targeted for degradation during senescence and liver aging, and loss of SIRT7 plays a crucial role in both the initiation of senescence and the control of metabolic and inflammatory genes in aging livers. (**Fig. 1** and **Fig. 2**). We discover interaction of SIRT7 with HMG Box containing transcription factor NUCKS1 in proliferating cells and deacetylations of NUCKS1; SIRT7 loss in senescence leads to NUCKS1 recruitment onto chromatin (**Fig. 3** and **Fig. 4**). Further, depletion of NUCKS1 delays senescence and diminishes expression of SASP genes mediated by senescence-associated transcription factors, NFkB and CEBPβ (**Fig. 5**). We translated these findings to mouse aging, and show that aged livers have significantly increased NUCKS1 on chromatin, further magnified in SIRT7 KO liver (**Fig. 6**). Analysis of NUCKS1 ChIP-seq data in hepatocytes identified a cohort of SIRT7 and aging DEGs that overlap with NUCKS1 binding peaks, including NFkB associated transcription factor *Rela* (**Fig. 7**). Overall, we propose that post-transcriptional degradation of SIRT7 releases NUCKS1 to localize to the nucleus, driving senescence-associated gene expression, in a model that is notably independent of histones as substrates of SIRT7 (**Fig 8**).

**Figure 8:**
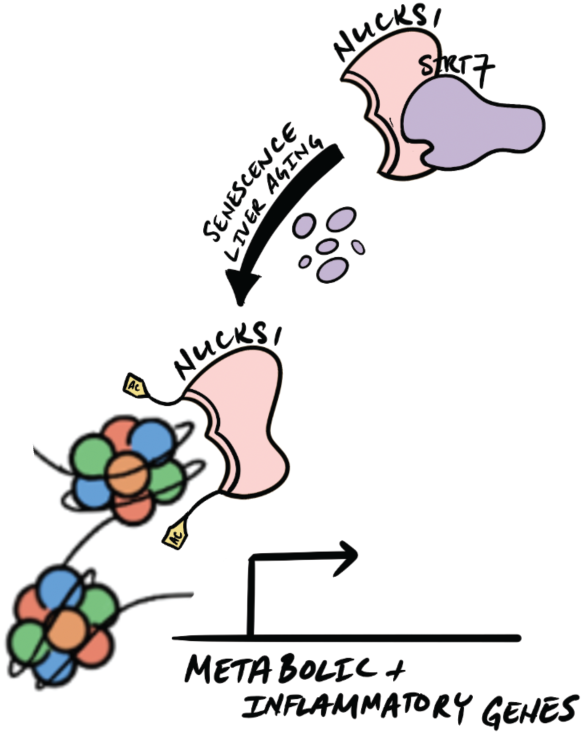
SIRT7 degradgation leads to NUCKS1 chromatin to mediate aging gene programs. Model of SIRT7 mediated regulation of NUCKS1 during aging and senescence. SIRT7 is targeted for degradation during senescence and aging leading to NUCKS1 acetylation and its recruitment onto chromatin. NUCKS1 mediates gene expression of key metabolic and inflammatory genes associated with aging.

The Sirtuin protein deacetylase family is strongly implicated in aging^4,5^. While SIRT7 is widely expressed across many tissues and cell types, we find the loss of SIRT7 protein to be specifically associated with liver aging compared to other tissues (**Fig. 2** and **Supp Fig. 2**). Similar observations were observed in rat where both SIRT6 and SIRT7 levels were reduced levels in aged livers^46^. SIRT7 transcripts are lower in hematopoietic stem cell aging resulting in disrupted metabolism through increased mitochondrial protein folding stress response and reduced regenerative potential^66^. SIRT1 protein is similarly reduced during senescence, however, SIRT1 degradation is mediated through the nuclear autophagy pathway^8^. These contrasting modes of degradation highlight the varying mechanisms of sirtuin regulation during stress. Further, SIRT1 is reduced in aging spleen and testes^8^; however, we observed no difference in SIRT1 in liver aging. These findings underline tissue-specific regulation of sirtuin proteins during aging and that selective upregulation of sirtuin proteins may contribute to healthy aging. We similarly showed that upregulation of SIRT7 during cellular senescence ameliorated characteristics of senescence and senescence-like SASP expression (**Fig. 1**), supporting the rationale of targeting SIRT7 to promote healthier aging. Relatedly, tissue-specific expression of Sirt7 within the vascular endothelium of Hutchinson-Gilford progeria syndrome mice improved aging characteristics including extended lifespan and reduced inflammatory response^67^. Identification of Sirtuin activators have been challenging, however, targeting the degradation of SIRT7 could be a more selective way to limit deleterious effects of SIRT7 loss in aging. A previous study showed that a component of a SCF E3 ubiquitin-protein ligase complex, FBXO7, can target SIRT7 for ubiquitination^68^. Additionally, our proteomic analysis of SIRT7 interactors identified several ubiquitin E3 Ligases (**Fig. 3**). It will be of interest to determine whether these E3 ligases mediate SIRT7 degradation and the potential targeted to maintain SIRT7 levels during aging.

While several studies implicate SIRT7 loss with senescence and aging, downstream mechanisms vary. For example, SIRT7 loss leads to ribosomal DNA damage and expression of LINE elements through dysregulation of chromatin^13,15^. SIRT7 deacetylation of histones and chromatin regulation has been a key focus, however there is increasing evidence of SIRT7 interacting with transcription factors to elicit transcription programs. For example, SIRT7 suppresses transcription factor HMG box SOX9 in chondrocytes^69^ and SIRT7 regulates cellular localization of transcription factor HMGB1 during hepatocyte stellate cell activation^22^. Here, we identified interaction of SIRT7 with the HMG protein NUCKS1 (**Fig. 3C**, **Fig. 4A**), which has been associated with processes associated with senescence^25–28^. SIRT7-NUCKS1 interaction sequesters NUCKS1 from chromatin binding and loss of SIRT7 during both senescence and liver aging licenses NUCKS1 binding to chromatin (**Fig. 4D-F**, **Supp Fig. 4I**). Importantly, depletion of NUCKS1 leads to massive changes in transcription, including genes linked to senescence and aging, and reduces senescence phenotype (**Fig. 5**). Thus, our results underscore SIRT7’s substrate diversity beyond histones in regulation of gene expression, to alteration of transcription factor activity.

NUCKS1 is ubiquitously expressed across all tissues involved in cell cycle regulation, DNA damage, and metabolism^25–28,50,51^—processes implicated in senescence and aging, and thus pointing to SIRT7-NUCKS1 interaction contributing to SIRT7’s regulation of senescence and aging. NUCKS1 is heavily post-translationally modified and highly disordered, including phosphorylation by cyclin kinases during cell division^53,54^, and acetylation, previously without clear function^53^. Here we identified NUCKS1 acetylation during senescence, at several lysines within and surrounding the DNA binding domain (**Fig. 4F**). Interestingly, we found SIRT7 to regulate several of these lysine residues, however, several senescence-associated NUCKS1 acetylation sites were not mediated by SIRT7 suggesting other deacetylases involved in NUCKS1 regulation (**Fig 4F**). The altered function of NUCKS1 during senescence, involving increased NUCKS1 chromatin binding (**Fig. 4C-D** and **Fig. 4F**), and senescence gene regulation (**Fig. 5**), suggests potential impact of NUCKS1 acetylation.

There is limited understanding of transcription-related functions of NUCKS1 beyond binding to insulin genes to promote transcription^50^. We compared ChIP-seq NUCKS1 binding with liver histone modifications and chromatin accessibility, revealing NUCKS1 association with regulatory enhancers marked by H3K27ac and H3K4me1 (**Fig. 7A** and **Fig. 7B**). Interestingly, NUCKS1-bound regions gain accessibility (ATAC-seq peaks) during liver aging, suggesting NUCKS1 promotes chromatin opening (**Fig. 7C** and **Fig. 7D**). We found enhanced NUCKS1 binding to the chromatin compartment during liver aging and in SIRT7 KO livers (**Fig. 6**). This predicts that NUCKS1 binding may promote gene expression and, indeed, NUCKS binding peaks occur proximal to SIRT7-regulated genes (genes more highly expressed in SIRT7 KO compared to WT) (**Fig. 7E**).

The NFkB family of transcription factors, involved in inflammatory response and senescence, consist of multiple members, including NFKB1 and RELA^64^. NUCKS1 knockdown during senescence identified diminished expression of a cohort of NFKB1 and RELA target genes—predominately inflammatory genes (**Fig. 5**). These results support previous studies in corneal epithelial cell wound healing that NUCKS1 promotes NFkB activity and in inflammatory and LPS stimulation pathways^26^. We find NUCKS1 binding at the *Rela* promoter in hepatocytes and *Rela* transcript levels increase during liver aging and in SIRT7 KO livers (**Fig. 7K** and **Fig. 7L**). However, in IMR90 senescence we find RELA expression is unaltered upon NUCKS1 depletion (**Supp Fig. 5J**) thus suggesting alternative mechanisms of NFkB regulation in senescence compared to liver aging.

In summary, our study highlights a novel mechanism exerted by SIRT7 in senescence and aging via NUCKS1 regulation, to elicit transcriptional programs associated with senescence and liver aging. These findings provide a potential target in SIRT7-NUCKS1 interaction to limit the detrimental effect of senescent cells in aging.

## MATERIALS AND METHODS

### Cell Lines

IMR90 (Primary human lung fibroblast) and HEK293T (Human kidney cell line) were described previously. IMR90 and HEK293T cells were cultured in DMEM supplemented with 10% fetal bovine serum (FBS) and 100 units ml−1 penicillin and 100 µg ml−1 streptomycin (Invitrogen). IMR90 cells were cultured under physiological oxygen (3%). IMR90 cells with a population doubling of less than 40 were used for experiments, except for replicative senescence. Stable cell lines were made by retrovirus or lentivirus infection, as previously described. For oncogene induced senescence (OIS), IMR90 cells expressing ER:HRasV12 were derived and treated with 4-hydroxytamoxifen (Sigma-Aldrich Catalog# T176) to induce HRasv12 expression. Uninduced cells were used as proliferative control. For DNA-damage induced senescence, IMR90 cells were transiently treated with 100uM of etoposide (Sigma-Aldrich Catalog# E1383) for either 24 or 48 hours. Cells were replaced with untreated media and assayed at designated time points. For replicative senescence (RS), IMR90 cells undergo continuous passage and are harvested at designated passage doublings. For MG132 (Calbiochem #474790) treatment, doses of 0.125–0.5 μM were used for 48-h treatments. For the Lys05 (MedKoo Biosciences Catalog# 406969) treatment, doses of 2–5 μM were used for 48-h treatments. For SIRT7 inhibitor 97491 (MedChemExpress #HY-135899), doses of 1.25-20 μM.

### Mouse Lines

129sv SirT7−/− mice (Vazquez et al., EMBO J. 2016) were housed at the Comparative Medicine and Bioimage Centre of Catalonia. All experimental procedures were ethically approved by the Catalan Ethical Committee and adhered to both international and national guidelines. For the collection of liver samples, young and old mice were humanely euthanized through CO2 gas inhalation. Following euthanasia, a precise incision was made in the abdominal area, and livers were carefully extracted. Subsequently, the livers were minced into small pieces using a razor, transferred to Eppendorf tubes, and rapidly snap-frozen in liquid nitrogen. Samples were stored at -80°C until needed for subsequent experimentation.

### Immunoblot

IMR90 cells were lysed in buffer containing 20mM Tris (pH 7.5), 137mM NaCl, 1mM MgCl2, 1mM CaCl2, 1% NP-40, 10% glycerol and supplemented with Halt Protease Inhibitor Cocktail (Thermo #78430) and Benzonase (Millipore #70746-3). Cells were incubated at 4C shaking for at least 1 hour. Sodium dodecyl sulfate (SDS) was added to a final concentration of 1% and boiled for 10 minutes. Samples were centrifuged at 13000 RPM and supernatant (lysate) was quantified for protein concentration. Lysates were subjected to electrophoresis and transferred to a 0.2uM nitrocellulose membrane. The membrane was blocked with 5% Milk for at least 30 minutes shaking and rinsed 3 times using TBS supplemented with 0.1% Tween (TBST). The membrane was incubated overnight at 4C with primary antibodies in TBST in 5% Bovine Serum Albumin (BSA) and probed with horseradish peroxidase-conjugated secondary antibodies and developed with SuperSignal West Pico Plus chemiluminescent substrate (ThermoFIsher #34580) and imaged using an Amersham Imager 600. For tissue samples, approximately 20mg of tissue was diced and rinsed with DPBS. Tissues were lysed in a RIPA buffer (50mM Tris pH 7.5, 150mM NaCl, 0.1% SDS, 1% Sodium Deoxycholate, 1% Triton X-100). Cells were then sonicated 30 seconds followed by 30 seconds off using Bioruptor for a total of 5 minutes. SDS was added to a final concentration of !% and boiled for 10 minutes. Samples were centrifuged at 13000 RPM and supernatant (lysate) was quantified for protein concentration and was used for immunoblot as described above.

### Immunofluorescence

Cells were washed with 4% Paraformaldehyde diluted in PBS solution and incubated for 10 minutes. Cells were then permeabilized by 0.5% Triton diluted in PBS for an additional 10 minutes. Samples were incubated in a fresh PBS blocking solution (5% Normal Goat Serum, 0.2% Tween-20, 0.2% Fish Skin Gelatin) for at least 30 minutes. Cells were stained in primary antibody (see Antibody Table) overnight at 4C. Cells were washed and stained in secondary antibody (1:1000) for 30 minutes at room temperature. Cells were mounted and imaged on Nikon Eclipse Ti2 using Nikon NIS software. Colocalization analysis was performed using CellProfiler (Ref) using MeasureColocalization module.

### B-galactosidase staining

Staining was performed using Millipore Cellular Senescence Assay Kit (#KAA002) following the manufacturer’s instructions. Cells were imaged in brightfield and percent positive was quantified across at least three images.

### RNA Isolation

Harvested cells were washed and incubated with 1mL of Trizol and incubated for 5 minutes. 200uL of chloroform was added per sample and vortexed. Samples were centrifuged and clear supernatant was harvested in new Eppendorf containing 318uL of 100% ethanol. Samples were transferred RNEasy column (Qiagen #74106) and centrifuged. Column were washed with 500uL of RW1 buffer and was treated with DNAse (Qiagen #79254) for 30 minutes at room temperature. Column was washed with 500uL of RW1 followed by 500uL of RPE buffer and eluted with 30uL of RNase free water.

### Quantitative PCR

Isolated RNA was reverse transcribed using High-Capacity cDNA Reverse Transcription Kit (ThermoFisher #4368813). cDNA was diluted 1:10 in H20 and was used for quantitative PCR using Power SYBR Green PCR Master Mix (ThermoFisher #4368708) following manufacturer protocol and run on Applied Biosystem QuantStudio Real-Time PCR System using targeted primer and reference primer sets (See Primer Table).

### RNA-Sequencing

RNA-sequencing libraries were prepared NEBNext Poly(A) mRNA Magnetic Isolation Module kit (NEB Cat #) followed by NEBNext Ultra II Direction Prep Kit for Illumina (NEB Cat #). Library sizes was measured using Bioanalyzer and pooled libraries concentration was calculated using NEBNext Lib Quant Kit (NEB Cat#). Libraries were pooled and sequenced on an Illumina NextSeq 550 using paired-end sequencing of 42 bases per read.

### RNA-Sequencing data analysis

SIRT7 knockout mouse liver RNA sequencing data were aligned to the mouse reference genome assembly mm9, and IMR90 NUCKS1 KD RNA sequencing data were aligned to human reference genome assembly hg19, using STAR. Reads for each gene with RefSeq annotations were counted using htseq-counts. Counts were normalized, and significant differences were calculated with DESeq2. Volcano plots of differential genes was visualized through R-package EnhancedVolcano. Gene ontology pathway analysis was performed through ConsensusPathDB (Ref).

### ChIP-Sequencing data analysis

ChIP dataset (See Dataset Table) were aligned to mm9 using bowtie2. After removing mitochondrial reads, PCR duplicates were removed using Picard. Peaks were called using MACS2 and either input (GSE58100) or young liver (GSE198189) was used as control. BigWig files were created using deepTools bamcoverage tool (Ref) and visualized by IGV (Ref). Metaplots and heatmaps were produced with deeptools (Ref). Peaks were annotated using ChIPpeakAnno R package (Ref).

### Cellular fractionation

Fractionation of proliferative and senescent cells was adapted from DeCaprio et al. (Ref) with minor modifications. All extraction buffers were supplemented with Thermo Scientific Halt Protease Inhibitor Cocktails to a final concentration of 1X. Cells were harvested and washed using cold PBS and cytoplasmic fraction was isolated by adding five time the pellet volume of digitionin extraction buffer and incubated on ice for 10-15 minutes. Samples were spun down at 480xg for 10 minutes at 4°C and supernatant (cytoplasmic fraction) was collected. Pellet was washed with digitionin extraction to remove residual cytoplasmic proteins. Pellet was then resuspended with five time pellet original cell volume with Triton X-100 extraction buffer and incubated at 4C rotating for 1 hour. Sample was centrifuged down at 8000xg for 10 minutes at 4°C. Supernatant (organelle fraction) was collected and pellet was washed with 1mL of Triton X-100 extraction buffer. Pelleted nuclei was lysed using 2.5 times original cell volume with Tween/DOC extraction buffer and incubated at 4°C rotating for 1 hour. Cells were centrifuged at 10000xg for 10 minutest at 4°C and supernatant (nuclear fraction) was collected. Pellet was washed with Tween/DOC extraction buffer to remove residual nuclear protein. Pellet was resuspended with 2.5 times original cell volume with Tween/DOC extraction buffer supplemented with MgCl2 (1mM final) and Benzonase and incubate at 4°C rotating. Samples were centrifuge at max speed (13000 RPM) for 10 minutes at 4°C and supernatant (chromatin fraction) was collected. Before downstream analysis, fractionation was validated via immunoblot using a cytoplasmic, nuclear, and chromatin marker (See Antibody Table).

### Tissue fractionation

Fractionation of liver tissue was performed using Thermo Scientific Subcellular Protein Fractionation Kit for Tissues (Catalog number: 87790) using the manufacturer’s instructions with slight modifications. ∼50mg of input liver tissue was used for fractionation.

### Mass spectrometry analysis for fractionated samples

Samples were analyzed in 6 biological replicates per caste with nanoLC-MS/MS. Peptides were separated using an UltiMate3000 (Dionex) HPLC system (Thermo Fisher Scientific, San Jose, CA, USA) using a 75 μm ID fused capillary pulled in-house and packed with 2.4 μm ReproSil-Pur C18 beads to 20 cm. The HPLC gradient was 0%–35% solvent B (A = 0.1% formic acid, B = 95% acetonitrile, 0.1% formic acid) over 90 min and from 45% to 95% solvent B in 30 min at a flow rate of 300nl/min.

The QExactive HF (Thermo Fisher Scientific, San Jose, CA, USA) mass spectrometer was configured following a data-independent acquisition (DIA) chromatogram library method (cite encyclopedia). For wide-window data, full scan MS spectra (m/z 385-1015) were acquired with a resolution of 60,000, AGC target of 1e6; MS/MS spectra were acquired with 8 m/z staggered isolation windows (normalized collision energy of 27.5 and an AGC target of 1e6). The chromatogram library was collected in 6 gas-phase fractions (GPF) per fractionated compartment with DIA with full scan MS spectra of 110 m/z each with 4 m/z overlapping windows with the same resolution and AGC targets settings as the wide window data (see Searle at al encyclopedia cite for windowing schemes). Mass spectrometry data files were demultiplexed with MSConvert (citation). Chromatogram library was built from 6 gas-phase fractions using Walnut in EncyclopeDIA (fasta information here). Wide-window data was processed in EncyclopeDIA (citation) with chromatogram library from Walnut. Processed mass spectrometry data files were imported into Skyline (citation). For statistical analysis, Skyline data was exported to MSstats (citation), normalized by equalizing medians, and a 2-tailed t-test was performed (significant if adjusted < 0.05). For NUCKS quantification, DIA data from mouse liver was analyzed with a targeted approach taking the best 2 precursors and 3 transitions based on retention time and fragmentation pattern using Skyline (version 22.2)

### Immunoprecipitation (IP)

Harvested cells were lysed in a buffer containing 20mM Tris (pH 7.5), 137mM NaCl, 1mM MgCl2, 1mM CaCl2, 1% NP-40, 10% glycerol and supplemented with Halt Protease Inhibitor Cocktail (Thermo #78430) and Benzonase (Millipore #70746-3). Cells were incubated at 4C shaking for at least 1 hour. Samples were centrifuged at 13000 RPM at 4C and supernatant (lysate) was incubated overnight with either antibody-conjugated Dynabeeds (Life Technologies XXXX) at 4C rotating. Beads were washed using lysis buffer for a total of 5 washes. For IP-immunoblot, beads were incubated with 1X sample buffer and boiled for 10 minutes to elute proteins. Beads were bound using a magnetic stand and elutant was used for immunoblot. For IP-MS, beads were resuspended in 50mM ammonium bicarbonate with DTT at a final concentration of 10mM and incubated for 30 minutes at 55C. Beads were treated with iodoacetamide at a final concentration of 25mM and incubated for 45 minutes in the dark. Trypsin was added a final concentration of XXX and incubated at room temperature overnight. Beads were bound to a magnetic stand and elutant was transferred to new tube and speed vacuumed til dry. Samples were resuspended in 0.1% TFA and washed using C18 stage tip. Samples were speed vacuumed til dry and resuspended in 0.1% formic acid before being injected into the mass spectrometer.

### Mass spectrometry analysis for IP-MS of FLAG-SIRT7

Samples were analyzed with nanoLC-MS/MS. Peptides were separated using an UltiMate3000 (Dionex) HPLC system (Thermo Fisher Scientific, San Jose, CA, USA) using a 75 μm ID fused capillary pulled in-house and packed with 2.4 μm ReproSil-Pur C18 beads to 20 cm. The HPLC gradient was 0%–35% solvent B (A = 0.1% formic acid, B = 95% acetonitrile, 0.1% formic acid) over 20 min and from 45% to 95% solvent B in 40 min at a flow rate of 300nl/min. The QExactive HF (Thermo Fisher Scientific, San Jose, CA, USA) mass spectrometer was configured following a data-dependent acquisition (DDA) method. All solvents used in analysis of MS samples were LC-MS grade. Full MS scans from 300-1500 m/z were analyzed in the Orbitrap at 120,000 FWHM resolution and 5E5 AGC target value, for 50 ms maximum injection time. Ions were selected for MS2 analysis with an isolation window of 2 m/z, for a maximum injection time of 50 ms and a target AGC of 5E4. MS raw files were processed with Proteome Discoverer version 2.3 and MS spectra were searched against a target + reverse database with the SEQUEST search engine using H. sapiens FASTA databases (reviewed, canonical entries, downloaded 9/27/21). For global proteome samples, iBAQ quantification was performed on unique + razor peptides. Trypsin cleavage was specific with up to 2 missed cleavages allowed. Match between runs was enabled but restricted to matches within a single biological replicate by separation of replicates into independent searches. Match between runs parameters included a retention time alignment window of 20 min and a match time window of 0.7 min. False discovery rate (FDR) was set to 0.01.

### Identification of protein acetylation

FLAG tagged NUCKS1 was immunoprecipitated in proliferating, senescent or TSA+NAM treated IMR90 using FLAG antibody (See Supplementary Table XXXXX). IP samples were run on a 4-12% Bis-Tris gel and washed three times with water before staining with Gelcode Blue Stain Reagent (Thermo Fisher 24590) overnight rocking. The stained gel was destained overnight in water. NUCKS1 band was excised, diced, and washed with 100mM ammonium bicarbonate before destaining overnight in 50% acetonitrile in 100mM NH4CO3. Destained gel pieces were dried by adding acetonitrile to cover for 5 minutes shaking followed by speed vacuum for 10 minutes. Freshly made 10mM DTT was added to dehydrated gel pieces and incubated for 1 hour shaking at 56C to reduce pieces. 55mM of iodoacetamide in 50mM NH4CO3 was added and incubated in the dark for 45 minutes. Gel pieces were dehydrated by adding acetonitrile and incubated for 5 minutes followed by speed vacuuming for 30 minutes. Gels were rehydrated with trypsin at a concentration of 10ng/uL diluted in NH4CO3 and incubated overnight at room temperature. NUCKS1 peptide was extracted by rinsing gel with 50mM NH4CO3 for 45 minutes shaking and the supernatant was transferred into new tube. Gel pieces were dehydrated with 50uL of 50% acetonitrile with 5% acetic acid and incubated at room temperature shaking for 15 minutes and the supernatant was collected. Samples was rehydrated with 50mM NH4CO3 for 15 minutes followed by dehydration with 100% acetonitrile for 15 minutes. Supernatant was collected and pooled with the previous collection. A final round of rehydration and dehydration was performed for a final extraction of NUCKS1 peptides. Pooled samples were speed vacuumed until dry and resuspended in 0.1% TFA for stage tipping.

### Statistical analysis

For two-sample comparisons of means, a paired Student’s t test was used, and significance was established by a p value ≤0.05. To identify differential expressed genes, DESeq2 analysis was perform with and significant genes was identified with a p-value ≤0.05. To assess differential peaks, MACS2 analysis was performed, and significance peaks were determined by a cutoff of q-value ≤0.01. To assess the enrichment of ChIPseq binding at various peak sets, the R package phyper was used to perform hypergeometric tests, and significance was established by a P value ≤0.05.

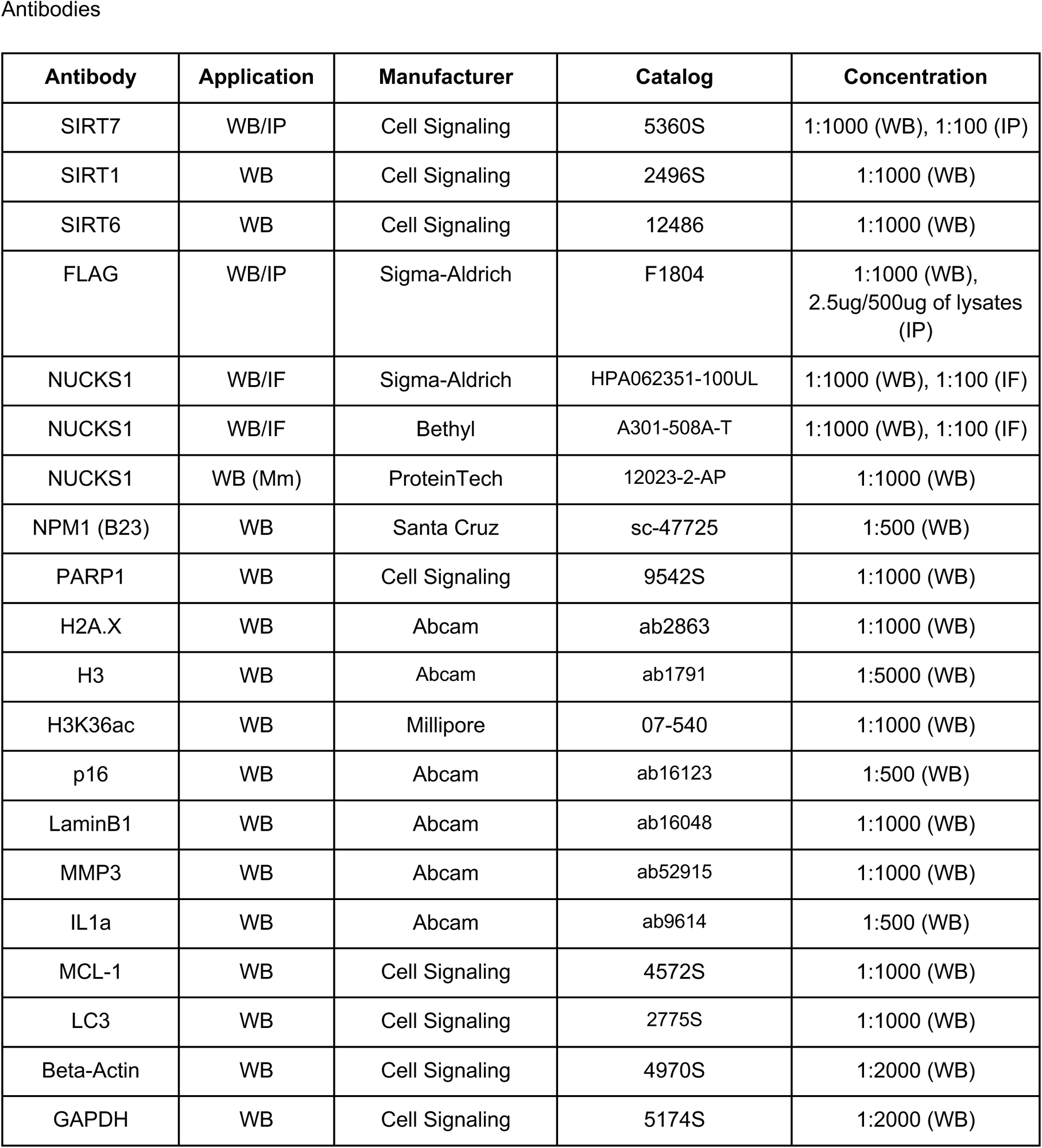
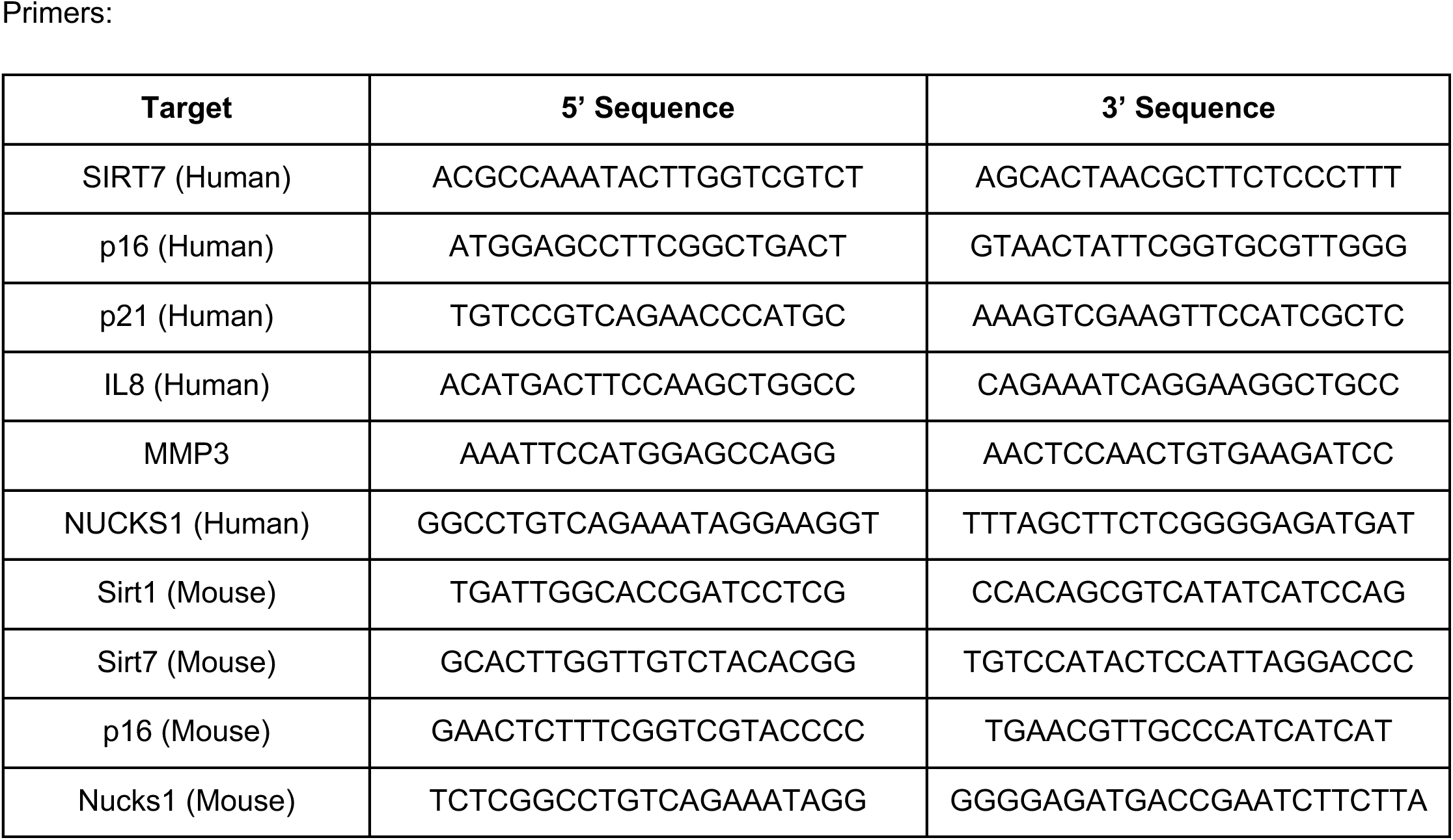
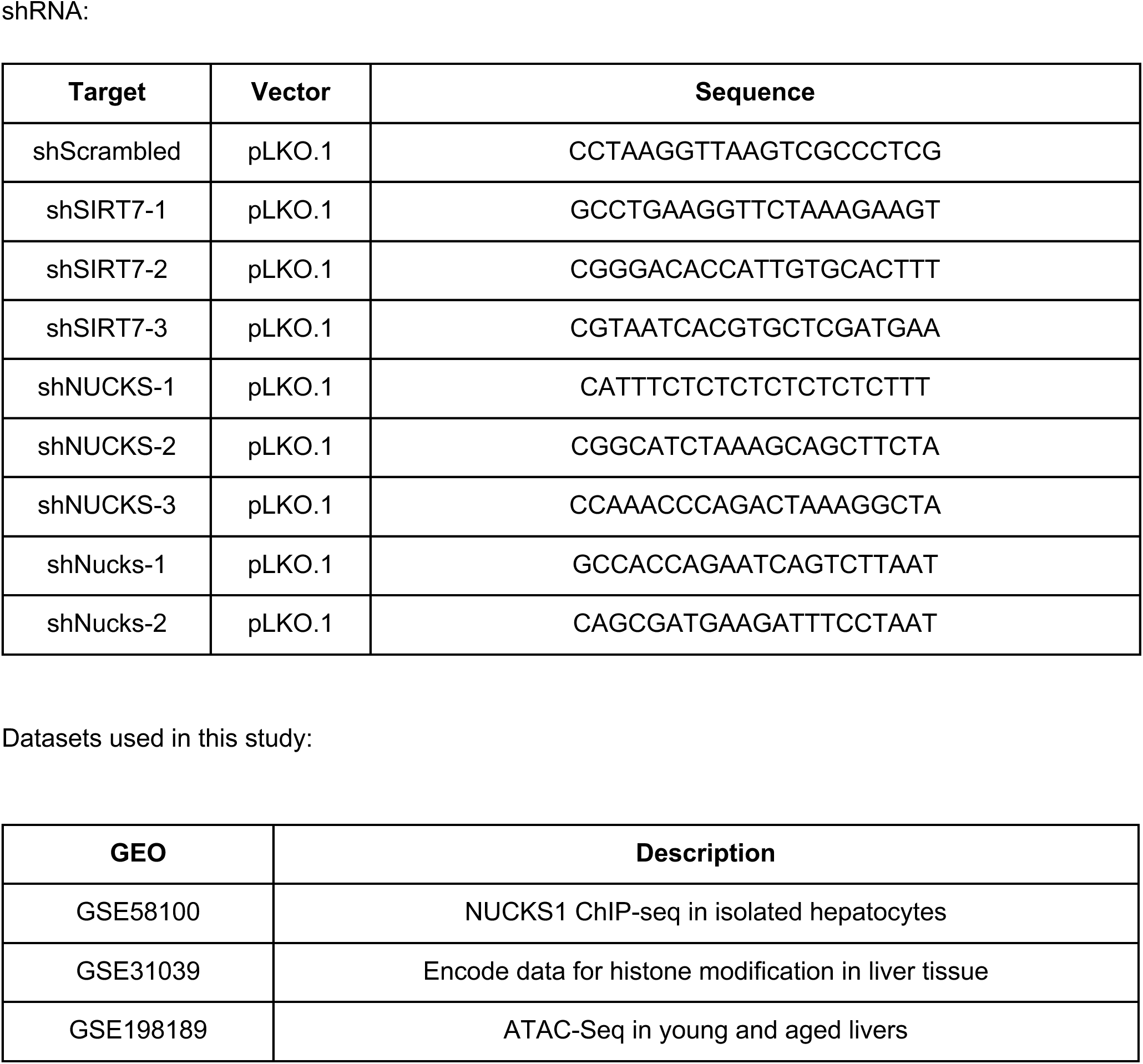

## Acknowledgements

We thank members of the Berger Lab (Charly Good) for help in editing the manuscript. We thank Brianna Aguilar and Zoe Moosbrugger for their help with cell culture. We thank Kate Alexander for assistance on imaging and image analysis. This work was supported by NIH training fellowships 1F32AG074641-01 (K.A.T.), NIA P01 AG 031862 (S.L.B.), P01AG047200 (B.A.G) and R01HD106051 (B.A.G).

---

**Supplementary Figure 1:**
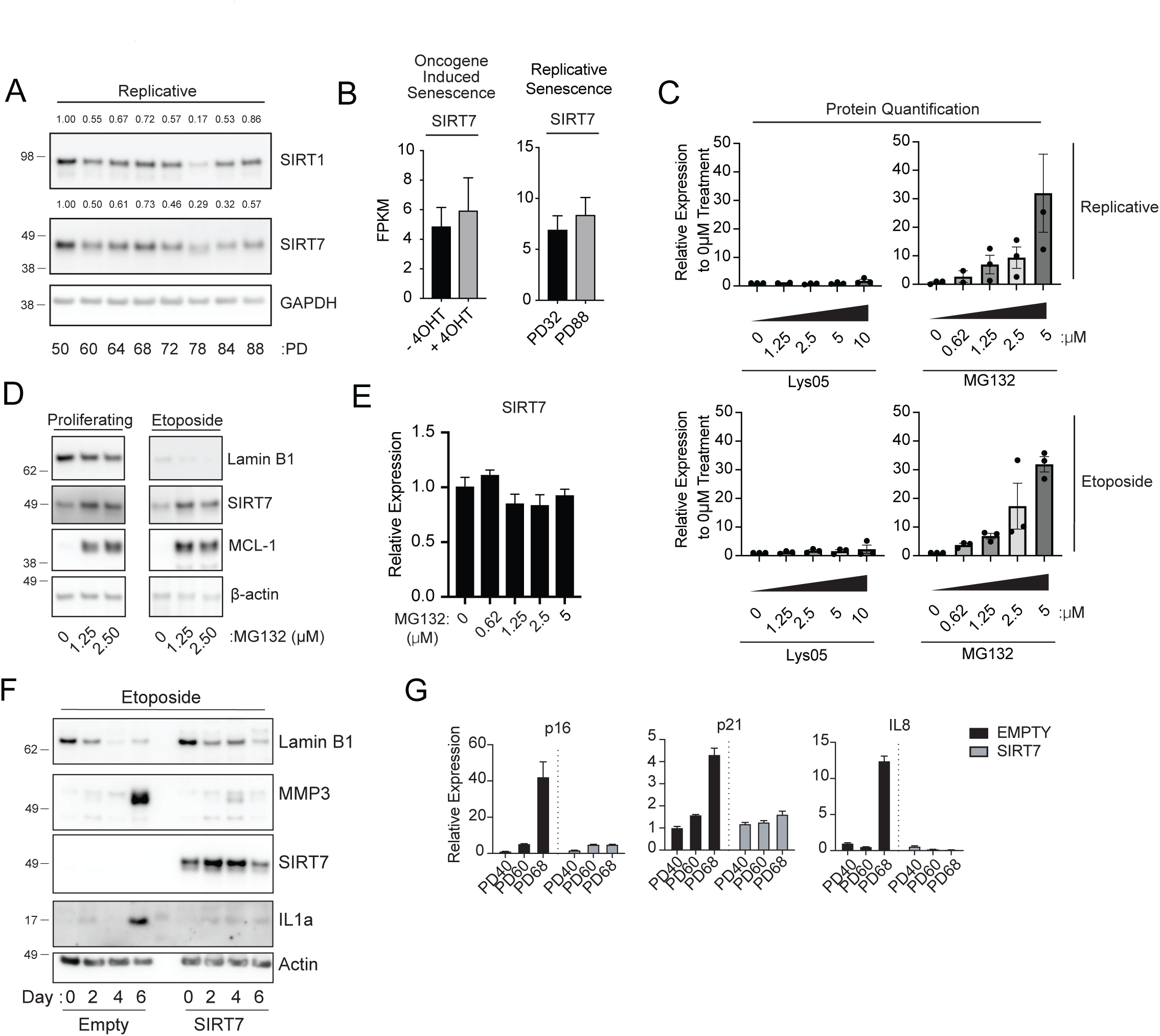
A. Immunoblot of SIRT1 and SIRT7 during replicative senescence. Cells were harvested at multiple population doublings (PD) until visual growth arrest (PD88). GAPDH is used as a loading control. B. Expression (Fragments Per Kilobase of transcript per Million) of SIRT7 from publicly available RNA-sequencing data in oncogene induced (left) or replicative (right) senescence. Error bars represent standard deviation of 3 independent experiments. C. Quantification of SIRT7 upon increasing dosage of either Lys05 or MG132 in either either replicative(Top) or etoposide induced senescent cells. Intensity was quantified by adjusting to loading control and normalized to 0μM. Bars represent average across three biological replicates. D. RT-PCR of SIRT7 in etoposide induced senescence treated with increasing concentration of proteasome inhibitor MG132. Values represent standard deviation of three technical replicates and normalized to Day 0. E. Immunoblot of proliferative (left) or etoposide induced senescent (right) cells treated with increasing proteasome inhibitor MG132. Lamin B1 depletion is shown as a marker of senescence and MCL-1 is used to quantify inhibition of protease activity. β-actin is used as a loading control. F. Immunoblot of IMR90 fibroblast expressing either FLAG-Empty or FLAG-SIRT7 during etoposide induced senescence. Cells were treated with etoposide for 24 hours and recovered for an additional 4 days. Cells were harvested at multiple timepoints and evaluated for senescence markers and SASP genes. G. RT-PCR of SASP genes MMP3 (Top) and IL8 (Bottom) of IMR90 cells overexpressing either empty or wildtype SIRT7 during replicative senescence. Cells were harvested at multiple population doublings (PD) until visual growth arrest (PD68) and evaluated for senescence markers and SASP genes. Error bars represent standard deviation of 3 independent experiments.

**Supplementary Figure 2:**
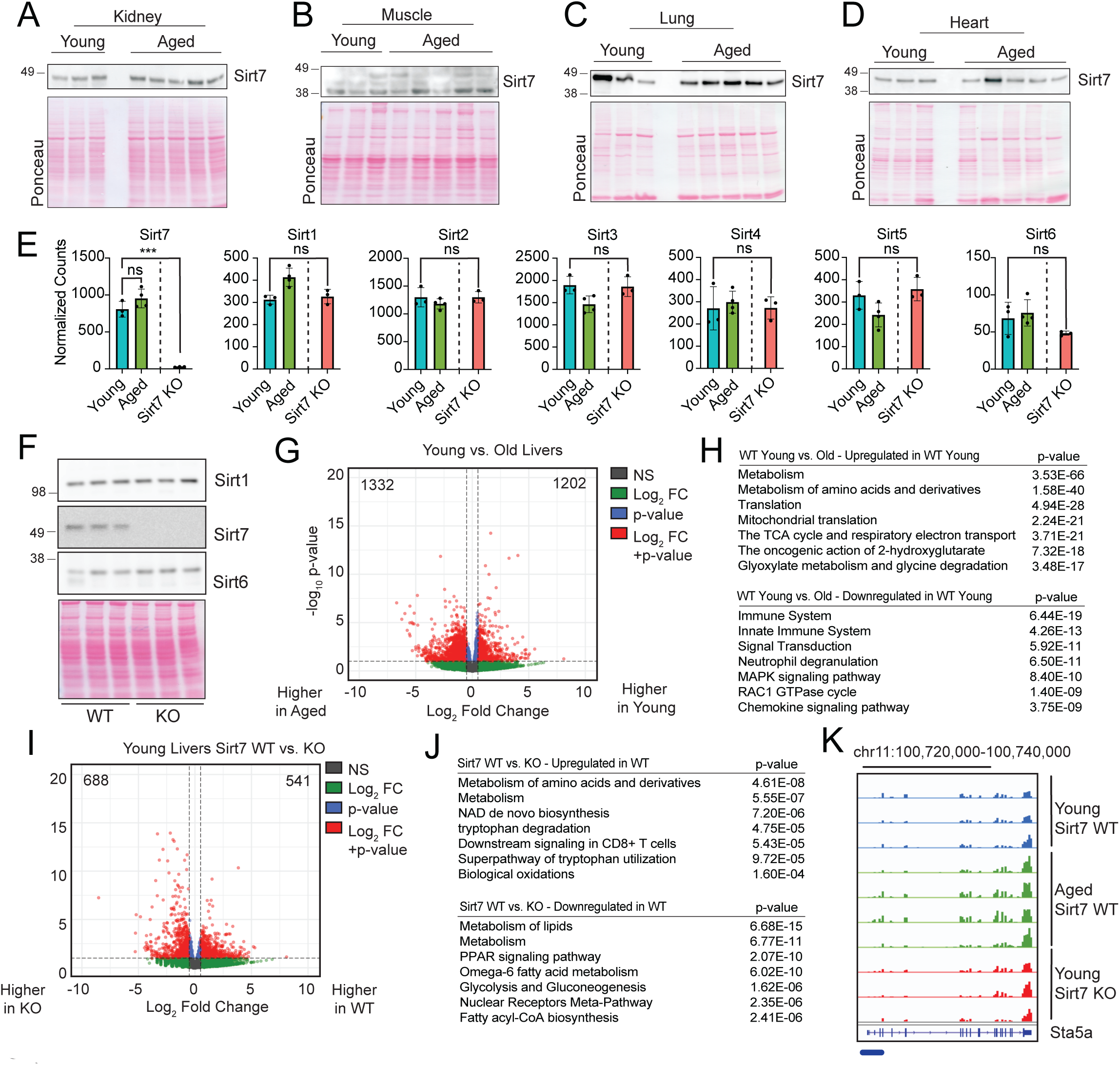
A. Immunoblot of SIRT7 in young and aged mouse kidney tissues. Ponceau is used as a loading control. B. Immunoblot of SIRT7 in young and aged mouse muscle tissues. Ponceau is used as a loading control. C. Immunoblot of SIRT7 in young and aged mouse lung tissues. Ponceau is used as a loading control. D. Immunoblot of SIRT7 in young and aged mouse heart tissues. Ponceau is used as a loading control. E. Normalized counts of sirtuin proteins for WT young (blue), aged (green), and young SIRT7 KO (red) liver samples. F. Immunoblot of nuclear sirtuin proteins in WT or Sirt7 KO liver samples. Ponceau is used as a loading control. G. Volcano plot of global transcriptomic analysis of Sirt7 wildtype young and aged mouse liver tissues. Colors correspond to the different cutoffs for p-value and fold change. Number of DEGs upregulated and downregulated are denoted in the top corners of the plot. H. Gene ontology (GO) for upregulated and downregulated DEGs between Sirt7 wildtype young and aged mouse liver tissues. I. Volcano plot of global transcriptomic analysis of wildtype young SIRT7 WT and KO liver tissues. Colors correspond to the different cutoffs for p-value and fold change. Number of DEGs upregulated and downregulated are denoted in the top corners of the plot. J. Gene ontology (GO) for upregulated (top) and downregulated (bottom) DEGs between SIRT7 WT and KO liver tissues. K. Gene tracks of RNA expression in young Sirt7 WT young (blue), WT aged (green), and Sirt7 KO young (red) livers at the Stat5alocus. Blue lines underneath tracks indicate promoter region.

**Supplementary Figure 3:**
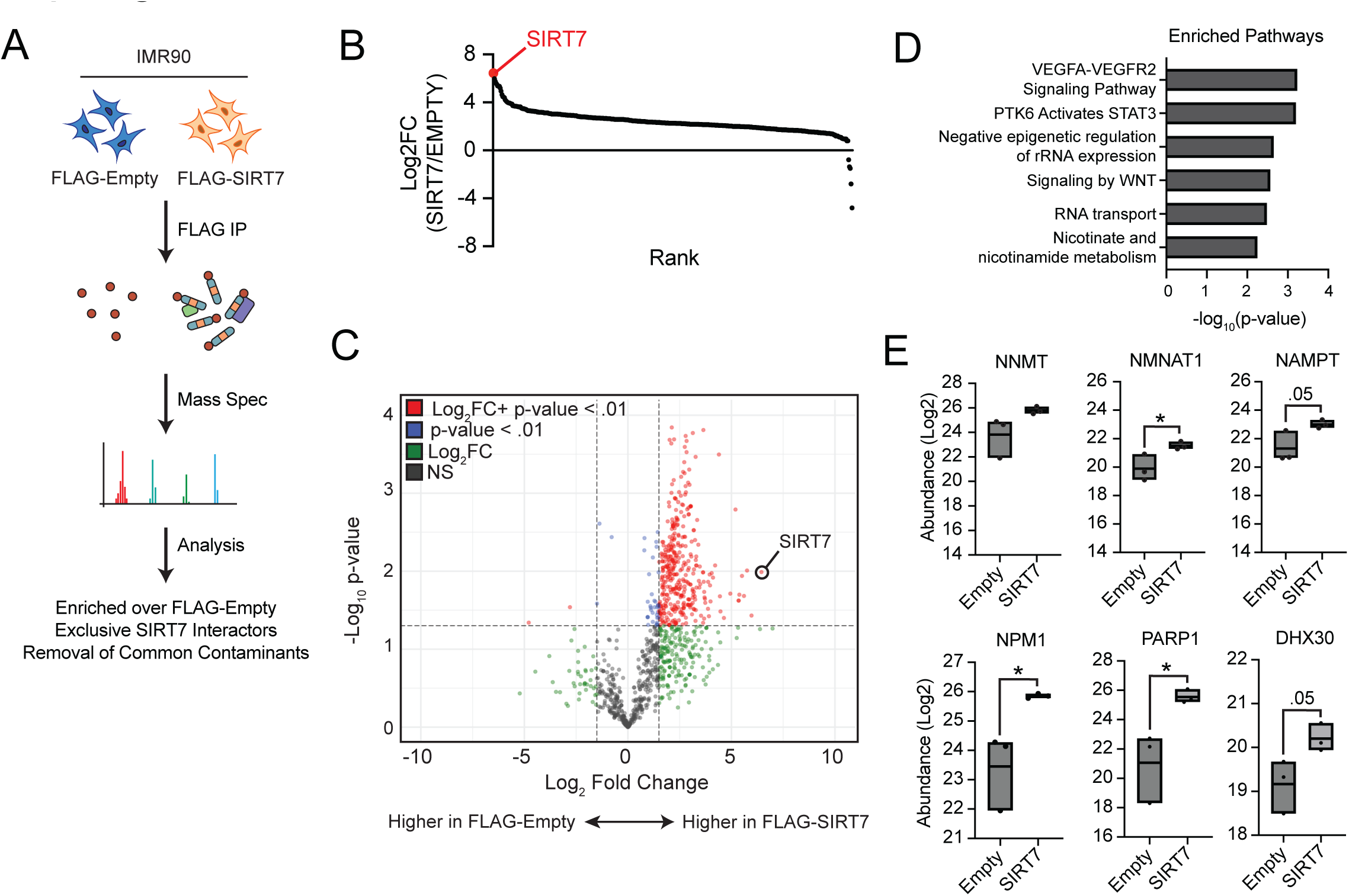
A. Schematic of identification of SIRT7 interactors in IMR90. FLAG-SIRT7 was immunoprecipitated in IMR90 cells followed by mass spectrometry. Interactors were identified by enrichment over FLAG-empty control followed by removal of common contaminants. B. Rank plot of significantly enriched proteins for either FLAG-SIRT7 or FLAG-Empty. C. Volcano plot of FLAG immunoprecipitation in IMR90 cells expressing FLAG-Empty or FLAG-SIRT7. Colors correspond to the different cutoffs for p-value and fold change. D. Gene ontology for the SIRT7 interacting proteins. E. Box plots of peptide abundance of selected proteins enriched in either FLAG-Empty or FLAG-SIRT7 immunoprecipitation. Interior line within box plot corresponds to mean across replicates. *** p < 0.001 * p < 0.05 = unpaired t-test.

**Supplementary Figure 4:**
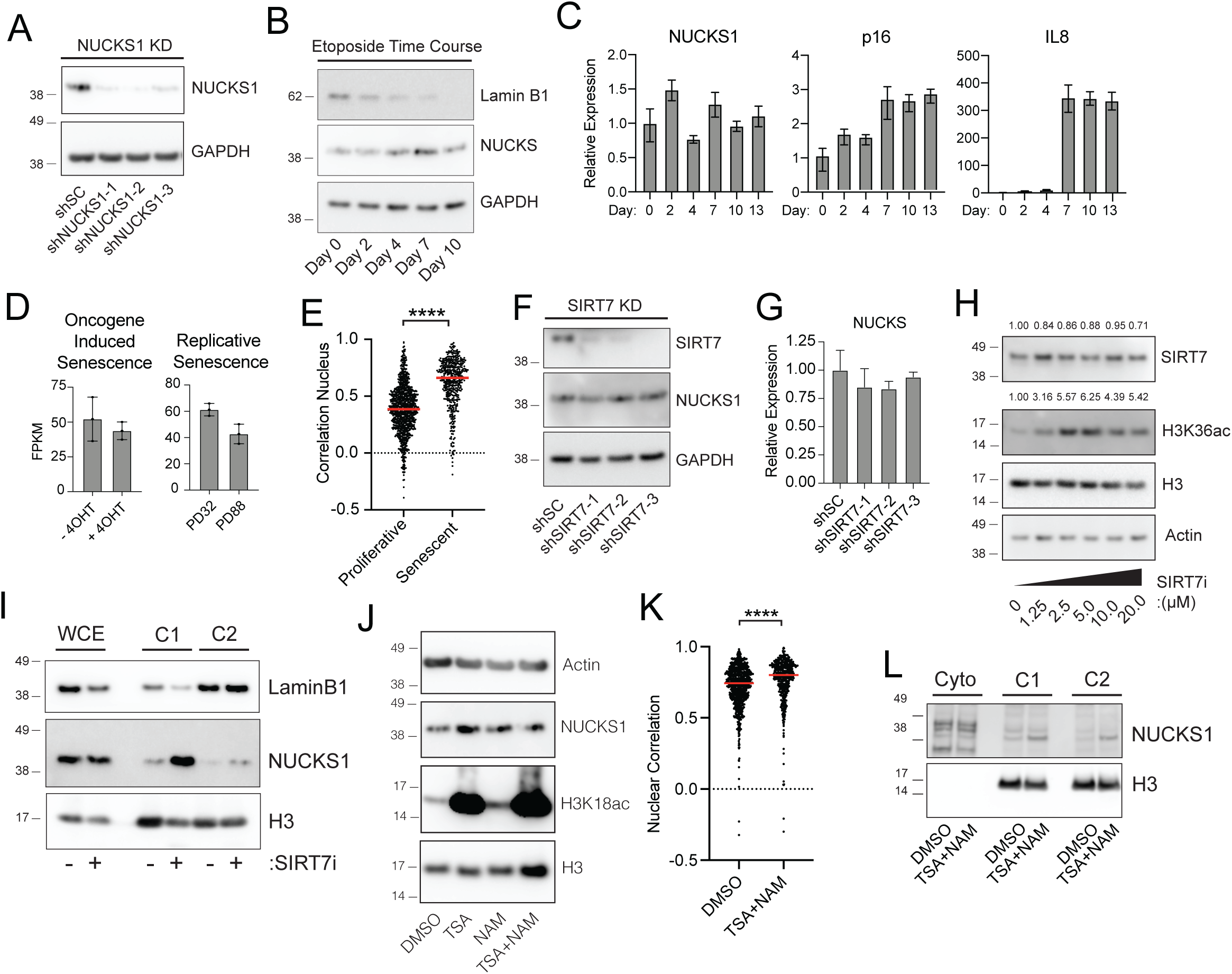
A. Immunoblot of NUCKS1 in IMR90 cells expressing shScrambled or three independent shRNA targeting NUCKS1. GAPDH is used as a loading control. B. Immunoblot of NUCKS1 during etoposide-induced senescence time course. Cells were treated with 100uM of etoposide for 24 hours and recovered in untreated media for an additional 8 days. Lamin B1 is used to evaluate senescence and GAPDH is used as a loading control. C. RT-PCR of NUCKS, senescence markers (p16) and SASP gene (IL8) during etoposide induced senescence. Cells were treated with 100uM of etoposide for 24 hours and recovered in untreated media for an additional 11 days. Values represent standard deviation of three technical replicates and normalized to Day 0. D. Expression (Fragments Per Kilobase of transcript per Million) of NUCKS1 from publicly available RNA-sequencing data in oncogene induced (left) or replicative (right) senescence. Error bars represent standard deviation of 3 independent experiments. E. Quantification of DAPI nuclear staining with NUCKS1 signal from immunofluorescence images with a second NUCKS1 antibody (Sigma). Each dot represents an individual nuclei with red horizontal bars indicating median value across all nuclei. **** p < 0.0001 = unpaired t-test. F. Immunoblot of NUCKS1 levels upon depletion of SIRT7 by shRNA. GAPDH is used as a loading control. G. RT-PCR of NUCKS upon depletion of SIRT7 by shRNA. Error bars represent standard deviation of 3 technical replicates. H. Immunoblot of H3K36ac upon increasing concentration of SIRT7 inhibitor (97491) for 48 hours. Intensity was quantified by adjusting to loading control (H3 for H3K36ac and Actin for SIRT7) and normalized to 0μM. I. Immunoblot of NUCKS1 in either whole cell extract (WCE) and two extractions of chromatin fractions (C1 and C2). Lamin B1 is used to indicate chromatin fraction and ponceau staining is used as a loading control. J. Immunoblot of NUCKS1 and H3K18ac upon treatment of either DMSO, TSA, NAM or TSA+NAM for 48 hours. H3 and Actin are used as loading controls. K. Quantification of DAPI nuclear staining with NUCKS1 signal from immunofluorescence images for IMR90 cells treated with either DMSO or TSA+NAM for 48 hours. Each dot represents an individual nuclei with red horizontal bars indicating median value across all nuclei. **** p < 0.0001 = unpaired t-test. L. Immunoblot of NUCKS1 in either whole cell extract (WCE) and two extractions of chromatin fractions (C1 and C2) in IMR90 cells treated with DMSO (Proliferating) or TSA+NAM for 48 hours. H3 are used as controls for chromatin fractionation.

**Supplementary Figure 5:**
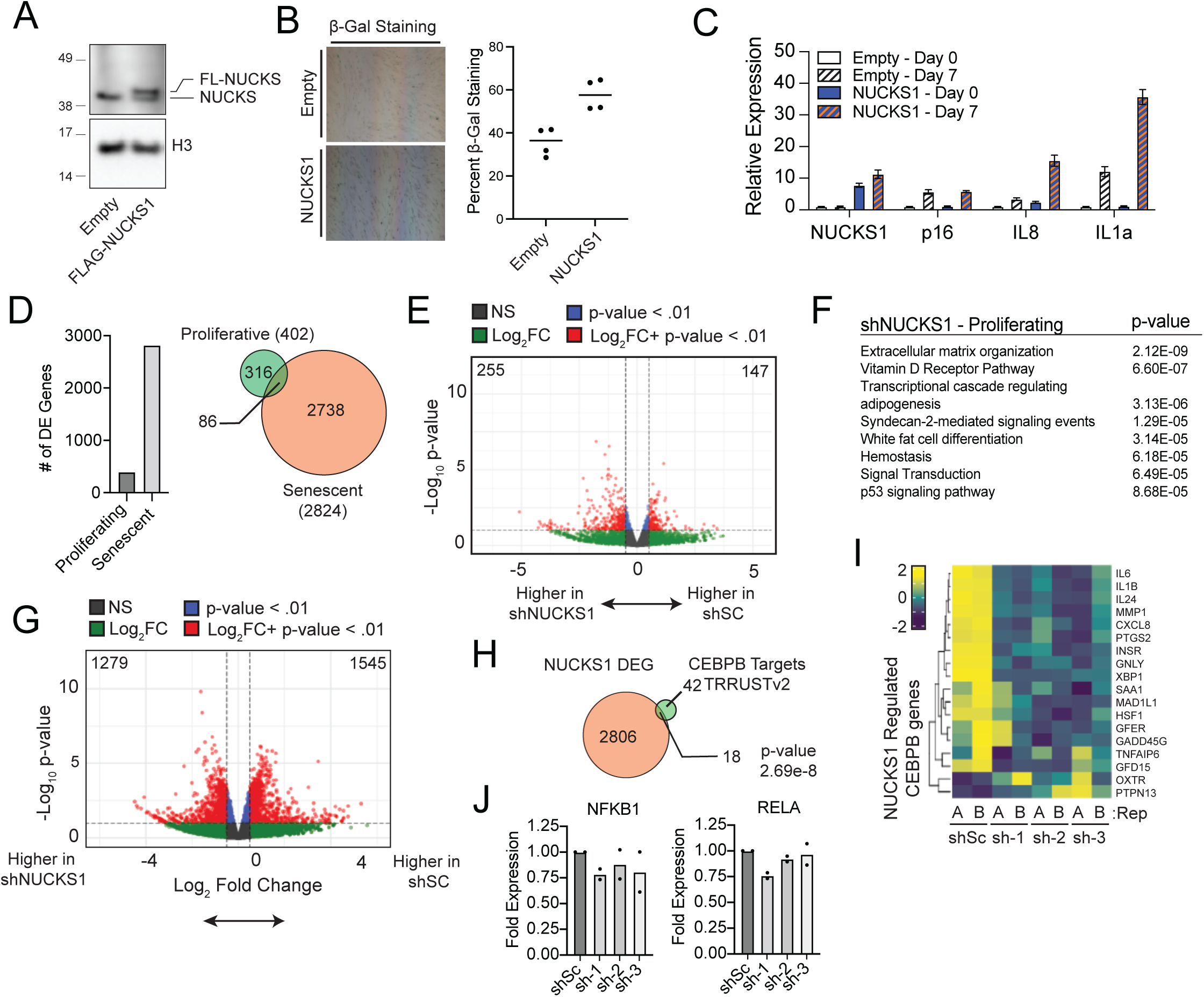
A. Immunoblot of NUCKS1 in IMR90 cells expressing FLAG-NUCKS1. H3 is used as a loading control. B. Left – representative images of b-galactosidase staining of IMR90 cells overexpressing FLAG tag alone (Empty) or FLAG-NUCKS1 (NUCKS1). Right - Quantification of percent B-galactosidase. Each dot represents a technical replicate with the solid bar indicating mean. C. RT-PCR of NUCKS1 (Left), senescence markers (p16) and SASP gene (IL8 and IL1a) during etoposide induced senescence for IMR90 expressing FLAG alone (Empty) or FLAG-NUCKS1 (NUCKS1). Cells were treated with 100uM of etoposide for 24 hours and recovered in untreated media for an additional 6 days. Values represent standard deviation of three technical replicates and normalized to Empty Day 0 sample. D. Left - Bar graph showing the number of NUCKS1 DEGs in proliferating and etoposide induced senescent cells. Right - Venn Diagram showing the overlap between NUCKS1 DEGs between proliferating (green) and senescent cells (orange). E. Volcano plot of global transcriptomic analysis between shScrambled and shNUCKS in proliferative IMR90 cells. Colors correspond to the different cutoffs for p-value and fold change. F. Gene ontology (GO) for DEGs for NUCKS1 depleted cells in proliferative cells. G. Volcano plot of global transcriptomic analysis between shScrambled and shNUCKS in etoposide induced senescent IMR90 cells. Colors correspond to the different cutoffs for p-value and fold change. H. Venn Diagram showing the overlap between NUCKS1 DEGs (orange) and CEBPB targets from TRRUST database (green). I. Heatmap of overlapped genes (18 genes from Supp Fig. 5G) of NUCKS1 DEG and CEBPB targets. Values represent Z-score across each replicate of shScrambled or shNUCKS1 depleted cells. J. Expression of NFKB subunits NFKB1 and RELA in etoposide induced senescent IMR90 cells. Values represent fold difference to shScrambeld control.

**Supplementary Figure 6:**
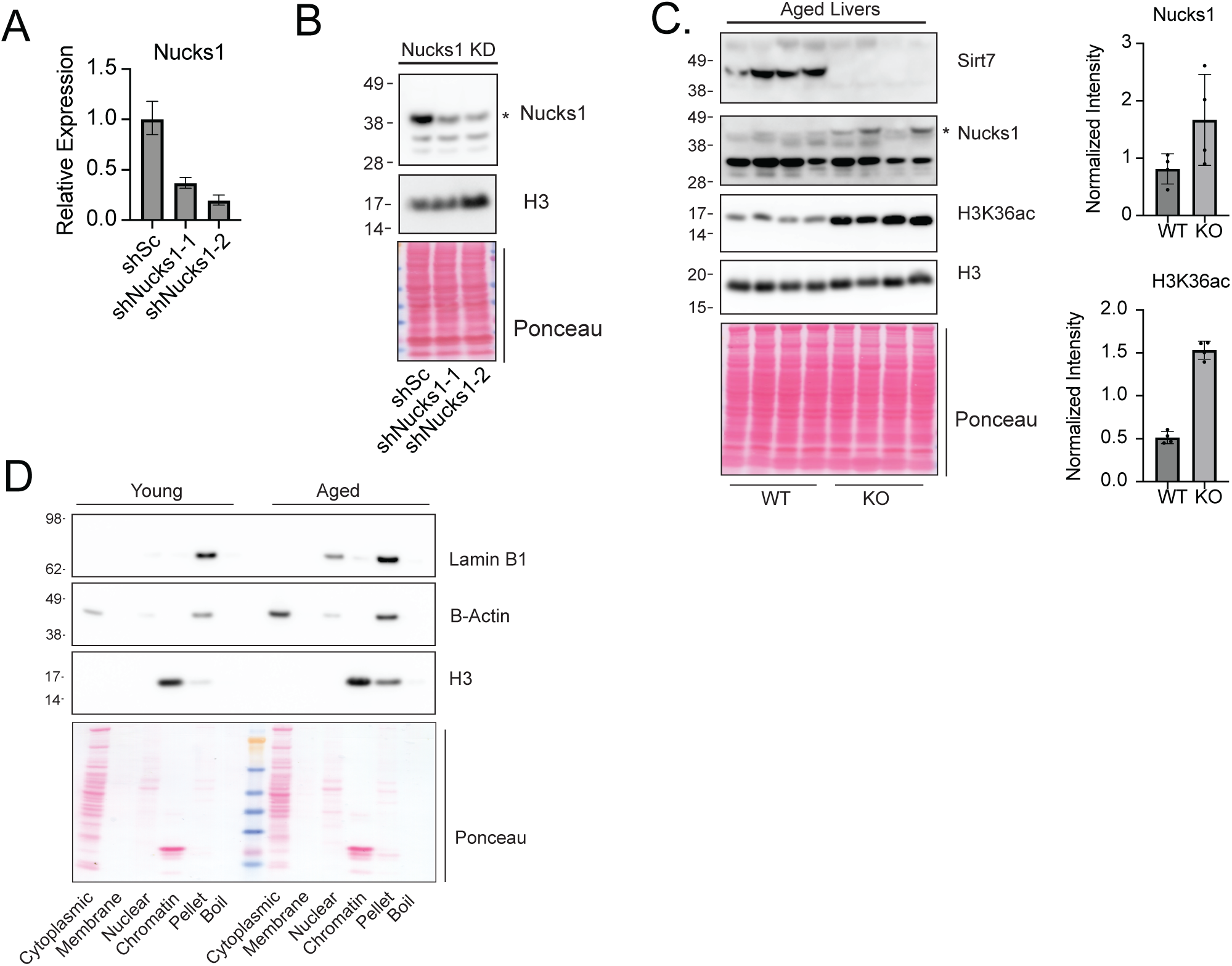
A. RT-PCR of Nucks1 in mouse EL4 cells treated with shRNA targeting scrambled control or Nucks1. B. Immunoblot of EL4 cells treated with shRNA targeting scrambled control or Nucks1 to validate antibody specificity (Nucks1 targeted band is denoted by *). H3 and ponceau is used as a loading control. C. Immunoblot of WT or KO Sirt7 in aged livers (Nucks1 targeted band is denoted by *). Ponceau staining is used as a loading control. Left - Quantification of Nucks1(Top) and H3K36ac (Bottom) in WT and KO Sirt7 Aged livers. Intensity was quantified by adjusting to H3 and normalized to median across all samples. D. Immunoblot of compartmental markers in different cellular fractions in young and aged livers. B-actin is used as a marker of cytoplasmic protein and H3 is used as a control for chromatin fractionation.

**Supplementary Figure 7:**
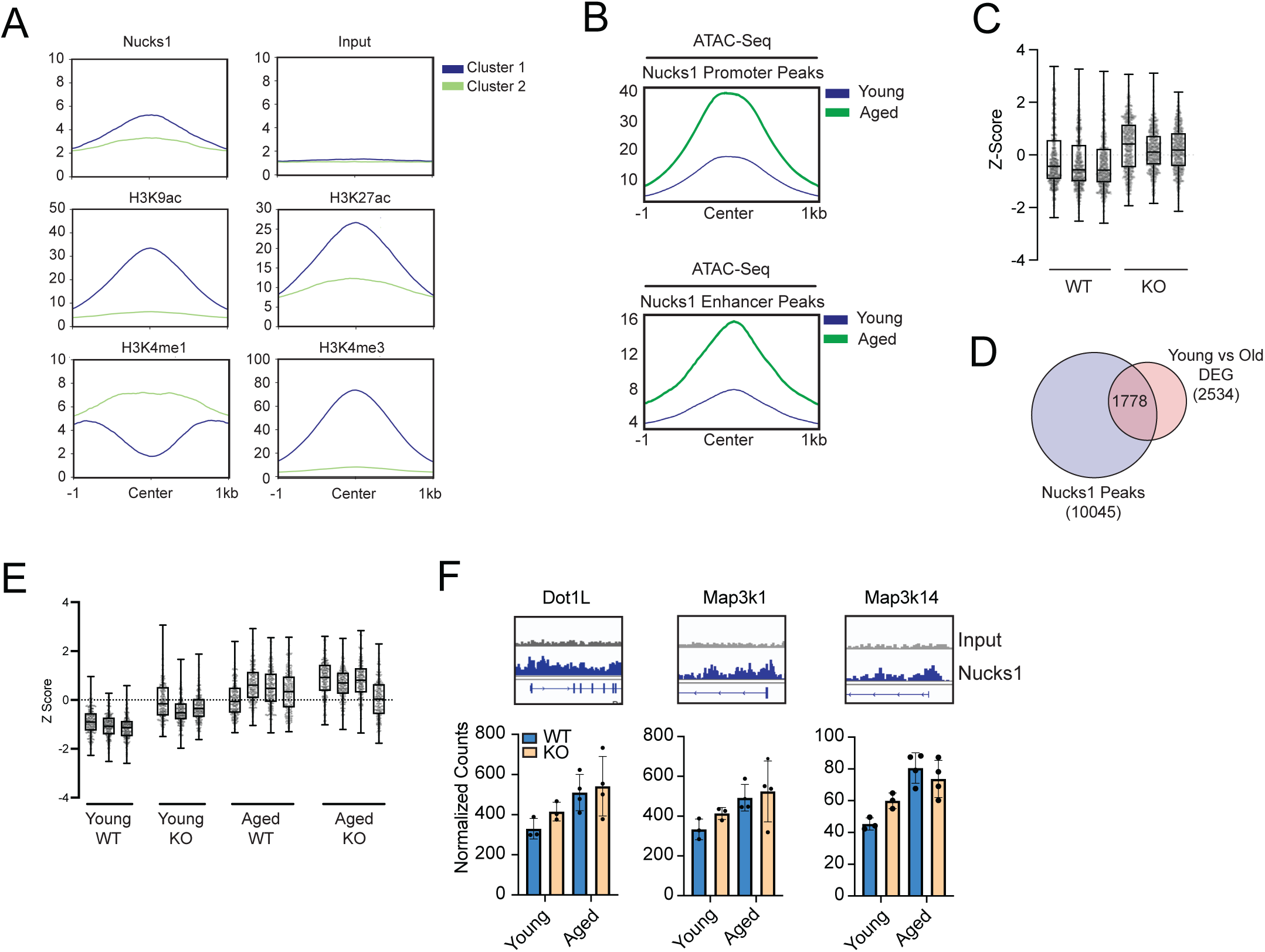
A. Metaplot of ChIP signal for Nucks1 called peaks for Nucks1, input (control), H3K9ac, H3K27ac, H3K4me1, and H3K4me3 in liver tissue for clusters 1 (blue) and cluster 2 (light green) defined in Fig 7A. B. Metaplot of ATAC-Seq signal at Nucks1 called peaks for promoters (cluster 1, Fig 7A) and enhancers (cluster 2, Fig 7A) in young (blue) and aged (light green) livers. C. Box plots of Z-score values from overlapped genes from Fig 7E for Sirt7 WT and KO young livers. D. Venn diagram of Nucks1 called peaks and differentially expressed genes (DEGs) in Sirt7 WT and KO young livers. E. Box plots of Z-score values from cluster 1 genes from Fig 7I for Sirt7 WT and KO in young and aged livers. F. Top - Gene tracks of ChIP-Seq signal for input (grey) and Nucks1 peaks (blue) for target genes. Bottom - Expression of target genes in Sirt7 WT and KO in young and aged livers.

